# Sequence determinants of intron-mediated enhancement learned from thousands of random introns

**DOI:** 10.1101/2024.10.29.620880

**Authors:** Emma J. K. Kowal, Yuta Sakai, Michael P. McGurk, Zoe J. Pasetsky, Christopher B. Burge

## Abstract

Spliceosomal introns are a ubiquitous feature of eukaryotic genes, whose presence often boosts the expression of their host gene, a phenomenon known as intron-mediated enhancement (IME). IME has been noted across diverse genes and organisms, but remains mysterious in many respects. For example, how does intron sequence affect the magnitude of IME? In this study, we performed a massively parallel reporter assay (MPRA) to assess the effect of varying intron sequence on gene expression in a high-throughput manner, in human cells, using tens of thousands of synthetic introns with natural splice sites and randomized internal sequence. We observe that most random introns splice efficiently and enhance gene expression as well as or better than fully natural introns. Nearly all introns stimulate gene expression ∼eight-fold above an intronless control, at both mRNA and protein levels, suggesting that the primary mechanism acts to increase mRNA levels. IME strength is positively associated with splicing efficiency and with the intronic content of poly-uridine stretches, which we confirm using reporter experiments. Together, this work elucidates sequence determinants of IME from tens of thousands of random introns, and confirms that enhancement of gene expression is a general property of splicing.

## Introduction

Despite appearing to be unnecessary for gene expression, introns are ubiquitous in eukaryotic genomes, with more than ten per protein-coding gene in humans on average (Koonin et al., 2013; Morales et al., 2022). The complex process of splicing required to remove introns from pre-mRNA is performed by the spliceosome, a dynamic macromolecular machine comprising many dozens of proteins and five small nuclear RNA-protein (snRNP) complexes (Maniatis and Reed, 1987; Papasaikas and Valcárcel, 2016). The spliceosome and other splicing regulatory proteins parse an extremely subtle and complex set of signals in the primary sequence of the pre-mRNA to identify introns and exons, signals which can be alternatively interpreted under different conditions to generate many isoforms from the same gene (Irimia and Roy, 2014; Papasaikas and Valcárcel, 2016). Each splicing reaction also results in the deposition of an exon junction complex (EJC), a large protein assembly which remains bound to the mRNA as it is exported from the nucleus into the cytoplasm (Le Hir et al., 2000; Saulière et al., 2012). Thus each intron, in the process of being spliced, orchestrates the interaction of the nascent transcript with a vast cohort of RNA-binding proteins (RBPs) and RNPs, some of which persist on the mRNA in its place long after its removal.

A curious aspect of intron splicing is that despite the extra time and effort required for their processing, introns generally enhance rather than inhibit gene expression (Fig. 1A). This was noticed very soon after the discovery of introns (Gruss et al., 1979; Hamer and Leder, 1979) and has since been described in diverse genes in a wide variety of plant, animal and fungal systems (reviewed in Shaul, 2017). Indeed, many promoters popularly used to drive expression of minigenes such as SV40, CMV, beta-globin, thymidine kinase (TK), Ubiquitin C (UbC), and EF1a commonly include a portion of 5ʹUTR with an intron that has been shown to boost expression significantly compared to the promoter alone. The magnitude of this intron-mediated enhancement (IME) can range from negligible to hundreds of times more mRNA and protein produced from a spliced versus intronless gene (Choi et al., 1991; Mascarenhas et al., 1990).

**Figure 1.**
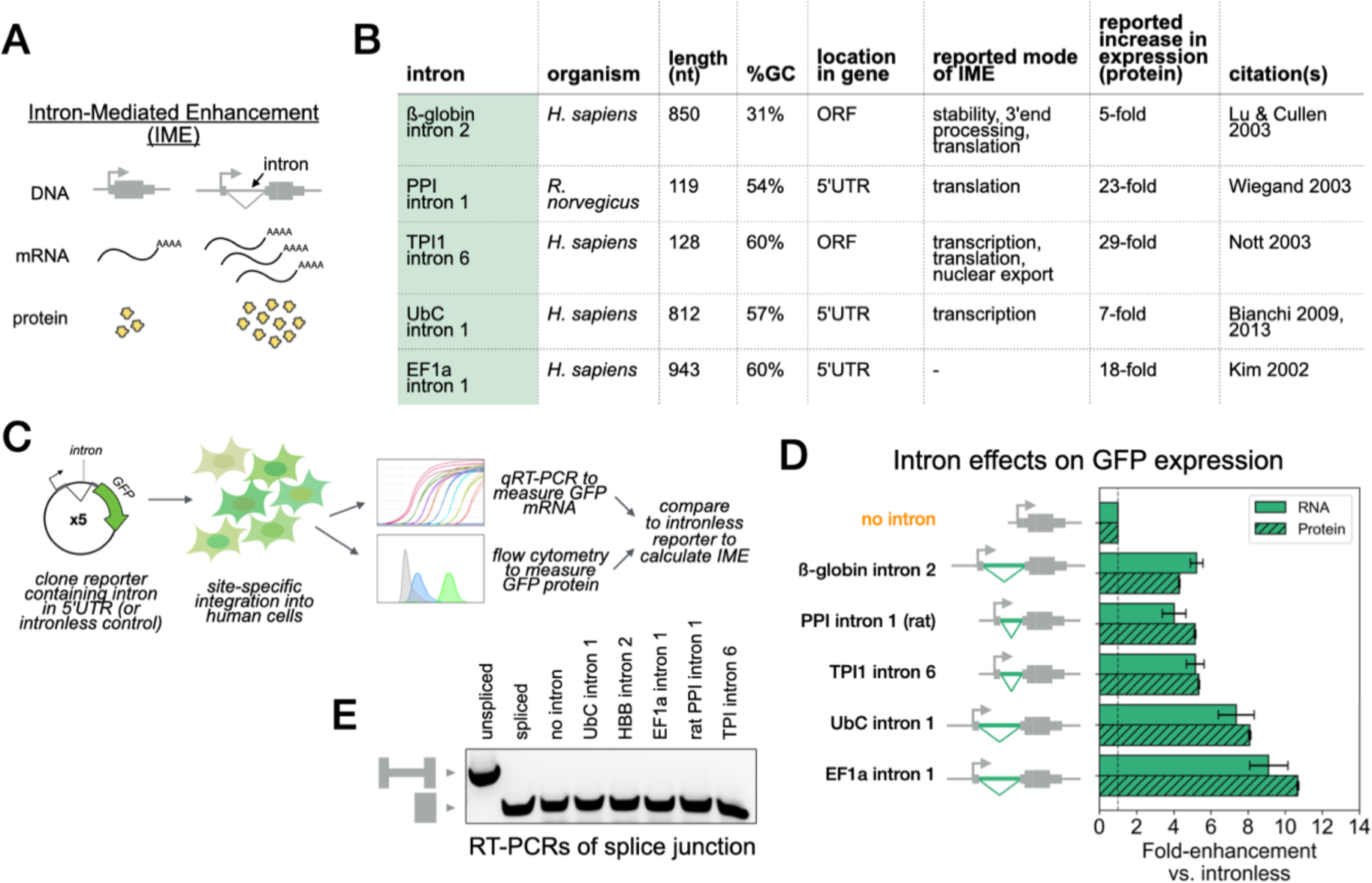
Different introns lead to different increases in gene expression. **A)** Schematic of intron-mediated enhancement (IME). **B)** Table of introns selected for testing in pilot experiment along with their reported modes and degrees of enhancement. **C)** Reporter experiment workflow. **D)** Pilot experiment showing effects on GFP reporter expression stimulated by insertion of different introns, using transgenic human HEK293T cells. Numbers are normalized to show fold change with respect to intronless control (dotted line). mRNA was measured from *n*=5 qRT-PCR replicates and protein from *n =* ∼20,000 cells using flow cytometry. Error bars denote standard error of the mean. **E)** RT-PCR of RNA from cell lines in D showing efficient splicing of all introns. Lanes 1 and 2 are controls for unspliced and spliced band size, PCR of plasmid DNA for UbC intron-containing and intronless reporters, respectively.

The mechanism of these effects is hypothesized to rely on each intron’s recruitment of *trans*-acting RNA and protein factors. The IME literature contains examples of introns that can positively impact nearly all stages of mRNA processing, from transcription and splicing through export, translation and decay, with individual introns often exerting multiple distinct effects (Chiou et al., 1991; Damgaard et al., 2008; Lu and Cullen, 2003; Millevoi et al., 2009; Nott et al., 2003; Valencia et al., 2008; Zhao and Hamilton, 2007; reviewed in Le Hir et al., 2003; Rose, 2018; Shaul, 2017). Certain general rules have emerged, such as that introns tend to enhance most strongly near the 5’ end of genes (Rose, 2004; Shaul, 2017; Dwyer et al., 2021), but otherwise any trend seems to have as many exceptions as examples.

One of the many questions about IME that remain unanswered: what is the effect of intron sequence on the magnitude of enhancement? It has been observed that different introns tested in an identical context can boost expression to varying extents, sometimes by up to an order of magnitude (Bartlett et al., 2009; Bourdon et al., 2001; Gallegos and Rose, 2015; Rose, 2004, 2002; Shaul, 2017; Yuan et al., 2013). For example, several introns, all well-spliced, exerted distinct effects in the GUS1 reporter in *Arabidopsis thaliana* (Rose, 2002). Subsequent development of a computational method to predict the magnitude of an intron’s IME in plants from its sequence – the IMEter – enabled the discovery of a motif, TTNGATYTG, which in reporter experiments transformed an intron with no IME into one that increased mRNA accumulation 24-fold and protein accumulation 40-fold relative to the intronless control (Rose et al., 2016). However, relatively little is known about the sequence determinants of IME in animals.

In this work, we sought to understand the effect of intron sequence variation on the strength of IME in human cells. We designed and executed a massively parallel reporter assay (MPRA) to test the expression of tens of thousands of unique introns, using high-efficiency and low-background (HILO) recombination-mediated cassette exchange (RMCE) technology (Khandelia et al., 2011). This allowed us to insert introns at single copy into a constant genetic context and to compare their expression at the mRNA and protein levels in a native, controlled genomic setting, revealing insights into the nature of IME.

## Materials and Methods

### Mammalian cells, plasmids and reagents

HEK293T A2 and HeLa A12 HILO-RMCE cells and plasmids pEM689 and pEM784 were kindly provided by Eugene Makeyev at Nanyang Technological University, Singapore (Khandelia et al., 2011). All oligos used in this research are listed in Supplementary Table S1 and were manufactured by Eton Bioscience or IDT. All plasmids used in this research are listed in Supplementary Table S2. All other reagents and their sources are listed in Supplementary Table S3.

### Cloning of individual intron reporters

All plasmids used in this study are adapted from pEM689. The CMV enhancer, chicken ß-actin promoter and chimeric intron from dTomato was replaced to minimal CMV promoter to create the workhorse backbone plasmid pEK1 for the library and individual intron reporters. EGFP gene with UbC promoter, UbC 5ʹUTR with intron, UbC 3ʹUTR and dTomato was inserted into pEK1 to construct the pilot construct pEK2 to insert the introns. The intron sequence in the UbC 5ʹUTR was variably replaced with alternative intron sequences or no intron. All assembled plasmids were transformed into *Mix & Go*! DH5 Alpha competent cells before plating on 2% LB-agar plates containing ampicillin (100µg/mL) and incubated at 37°C overnight to obtain colonies. The colonies were picked and cultured in 2mL LB media containing ampicillin (100µg/mL) and grown overnight to purify the plasmids using QIAprep Spin Miniprep kit. The complete sequence of the insert or the whole plasmid was confirmed by Sanger sequencing (performed by Eton Biosciences) or by full-plasmid sequencing (performed by Plasmidsaurus).

### Cell culture and generation of transgenic lines

Both HEK293T A2 and HeLa A12 HILO-RMCE cells were grown in DMEM, with high glucose and pyruvate, supplemented with 10% fetal bovine serum and 10% penicillin/streptomycin. Cells were passaged 1:10 every 2-3 days by washing with PBS and treated with 0.25% Trypsin following standard procedures and tested for Mycoplasma by PCR periodically. The intron reporter cassettes were integrated by following authors’ recommendations: briefly, we first plated 1.5×10^5^ cells per well on a 12-well plate coated with 0.1mg/mL Poly-d-Lysine. After culturing the cells for 24 hours in antibiotic-free media, the cells in each well were co-transfected using Lipofectamine 2000 with 500 ng of a reporter plasmid plus 0.5% wt/wt Cre recombinase vector pEM784 (Khandelia et al., 2011), and allowed them 24–48 hours before selection with 4–8µg/mL Puromycin for two weeks to ensure genomic integration. Cells were monitored throughout for surviving colony formation and wells were split and pooled before becoming overconfluent. At this stage aliquots of 0.5 to 5×10^6^ cells were frozen in media with 10% Dimethyl sulfoxide (DMSO) and later thawed again when necessary. For experiments with inducible promoters, cell lines with integrated dox-inducible reporters were treated with 0–16µg/mL Doxycycline in the cell culture media and allowed to grow for 24 hours before trypsinization and flow cytometry.

### RT-PCR and qRT-PCR

For each RNA extraction, 10^5^ – 10^6^ cells were washed once with PBS and either trypsinized and pelleted or lysed directly in the tissue culture well. RNA was subsequently extracted using either Zymo Quick RNA MiniPrep kit with in-situ DNase I treatment, or for higher throughput applications, with the Chemagic 360 RNA extraction protocol. Purified RNA was quantified by either Nanodrop or Synergy H1 plate reader and 300-1000ng RNA per sample was reverse transcribed using SuperScript IV VILO For RT-PCR to verify correct splicing, 2µL of each RT reaction was used as the template for a Polymerase-chain-reaction (PCR) using Phusion DNA polymerase PCR and exon-junction-spanning primers (UbC-splicecheck-3 and GFP-N-out). Products were separated and visualized using agarose gel electrophoresis with SYBRSAFE dye and relative band intensities were quantified using FIJI. For qRT-PCR, RT reactions and primers for genes of interested were arrayed in a 96-well plate on ice, mixed with PowerUp SYBR Green Master Mix, and then transferred in quadruplicate to a 384 well plate using the TECAN EVO. Plates were run on a QuantStudio 5 thermocycler (Thermo) using SYBR manufacturer’s recommendations for cycling protocol. Cq values (mean of 4 technical replicates) were downloaded and data was analyzed using custom Python scripts. Each plate contained a serial dilution with at minimum 5 samples for plotting a standard curve and extracting primer efficiencies. Relative quantities for each target gene were calculated between samples of interest and the intronless control samples by exponentiating their deltaCq values using target-specific amplification efficiencies. GFP and dTomato values were then normalized to the geometric mean of 3 control targets (*GAPDH, RPL27, SRP14*) before computing their log ratio, which was finally again normalized to the intronless log ratio.

### Flow cytometry

Flow cytometry experiments were performed by using either an LSR II HTS-1 or FACS Symphony A3 HTS-1 instrument (BD Biosciences) at the Koch Institute Flow Cytometry core. Cells intended for analysis were washed with PBS, trypsinized, pelleted, carefully resuspended, passed through a 35µm filter lid to dissociate any clumps (Corning) and transported on ice. Single live cells were identified by binning FSC-A/SSC-A, FSC-W/H and SSC-W/H, with binning performed by flow core staff. GFP was measured in the FITC channel (Laser: 488nm, Filter: 515/20) and dTomato in the PE channel (Laser: 561nm, Filter: 586/15).

### Cloning of random intron library

*See Fig. 2B for schematic of procedure. Full protocol available upon request*.

**Figure 2.**
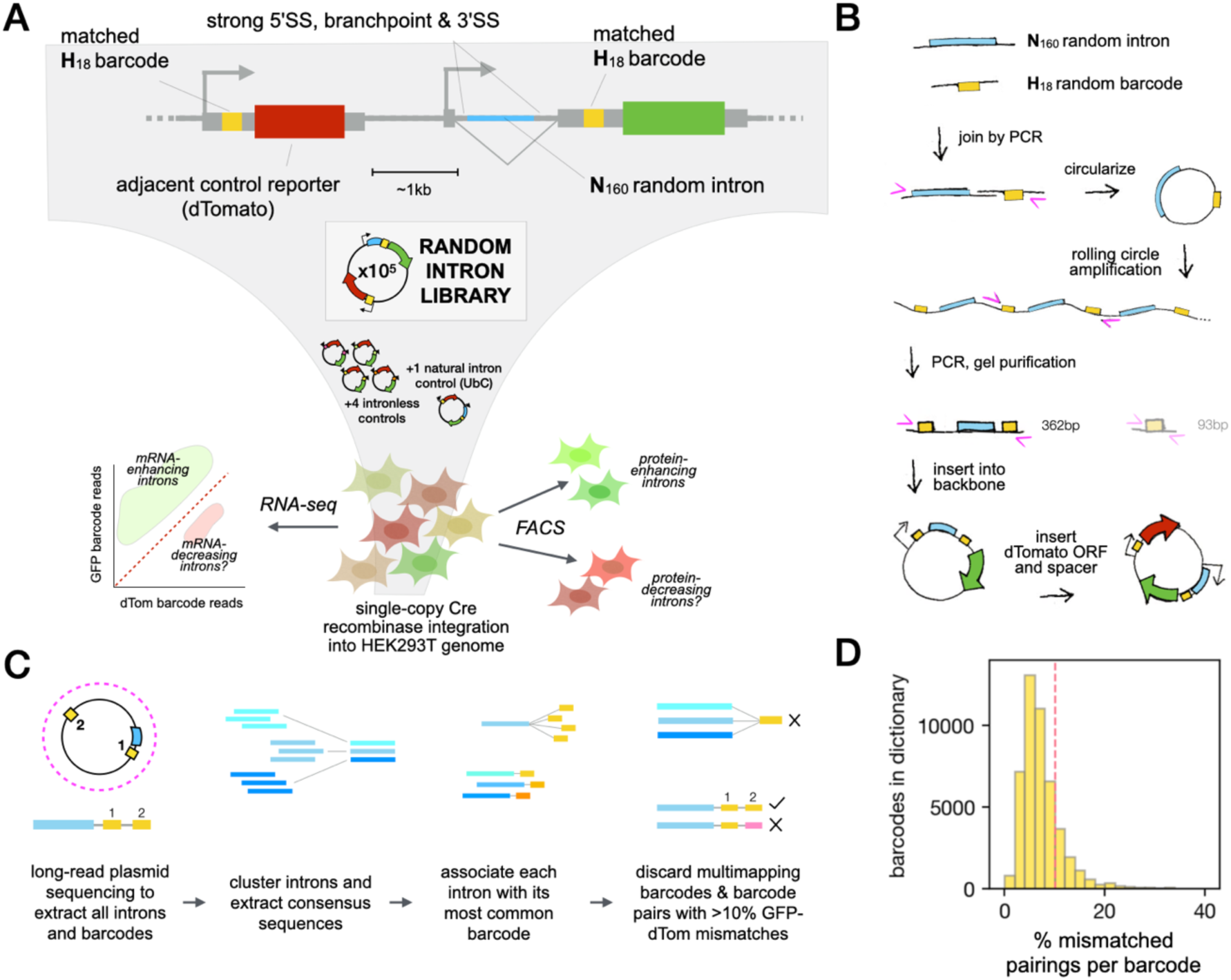
Development of a barcoded dual-reporter random intron library to measure IME effects of many introns in parallel. **A)** Schematic of reporter design and workflow for random intron screen. **B)** Cloning strategy for assembly of random library with random paired barcodes. H = A, C or T. **C)** Generation of barcode-to-intron map using reads from Oxford Nanopore long-read sequencing of library plasmid DNA. **D)** Distribution of mismatching rates between GFP and dTomato barcodes for all barcodes mapped to introns. Dotted line indicates cutoff for inclusion in further analyses (<10% mismatched).

### Preparation of acceptor plasmid for random intron library

Plasmid pLA (“Library Acceptor”) was constructed by Gibson assembly by introducing several changes to pEM689 and was used to insert the random intron library:

1. The GFP 5ʹUTR was changed into a hybrid between UbC and EF1a, extending the sequence between the intron 3ʹSS and the GFP start codon from 3nt to 32nt to allow for introduction of an 18nt barcode;
2. The dTomato promoter was changed from minCMV to UbC to match the GFP promoter;
3. Cut sites for the type IIS restriction enzymes BspMI and PsrI were introduced flanking the “dTomato Fragment” (from immediately upstream of the dTomato start codon to immediately upstream of the GFP start codon);
4. The bacterial EM7 promoter and Trimethoprim resistance gene were introduced between the dTomato and GFP, to ensure later re-insertion of the dTomato fragment.

Plasmid pLA was pre-digested with BspMI and gel purified to separate the dTom fragment from the GFP fragment. The dTom fragment was amplified by PCR using Phusion DNA polymerase, pre-digested with PsrI and gel purified again.

### PCR joining of random introns with random barcodes

Two degenerate oligonucleotides were used to create the library: EK148, a 190nt Intron Fragment, with 160nt internal random nucleotides flanked by 15nt constant sequence at either end for primer binding, and EK147, a 59nt Barcode Fragment, composed of 18xH nucleotides (A, T or C) flanked by 19nt and 22nt constant primer binding regions. 6ng of the intron fragment, theoretically ∼6×10^10^ unique sequences, was first amplified by 6 cycles of PCR in order to add the constant 5ʹ and 3ʹ splice site sequences, and to add a PsrI cut site upstream of the 5ʹSS. This PCR was performed in 50µL total volume with 2% DMSO using Phusion polymerase and GC buffer (NEB) with an annealing temperature of 52°C and extension time of 30 seconds. The product was purified with 1x SPRI beads (Agilent) and quantified by nanodrop. 1ng of this purified product was used as the template in a second PCR using a forward primer plus 2uL of the 100µM stock of barcode fragment, which acted as a reverse primer. This reaction was cycled 6 times to combine the two random regions and then again SPRI purified with 1x volume of beads. The subsequent 4 steps each took place with these SPRI beads remaining in the tube, in order to consume all of the reaction product and minimize sample loss from tube transfers, as in (Fisher et al., 2011). New reaction components were added directly to the eluted product and beads, and following the reaction, 20% polyethylene glycol (PEG) + 2.5 M NaCl buffer was added to achieve re-precipitation of DNA onto the same beads.

### Rolling Circle Amplification to generate introns flanked by duplicated barcodes

All steps in this section were performed with beads and followed by SPRI purification unless otherwise stated. The barcode-intron product was amplified by PCR once more and then digested with Lambda exonuclease to convert the double-stranded DNA into a single-stranded (ss) product for circularization. The purified ssDNA was treated with T4 Polynucleotide Kinase to phosphorylate free 5ʹ ends, then circularized overnight with Circligase II. This circularized product was eluted from SPRI beads in 4µL nuclease-free water, and 0.5µL was used as the template in rolling circle amplification (RCA) performed using the TempliPhi 100 Amplification Kit. The resulting linear concatemers were diluted 10× in nuclease-free water and sonicated in a Diagenode Bioruptor Plus sonication device at 4°C on low power for 5 cycles of 30sec on/90sec off. Sonicated RCA product was SPRI purified with 0.6× volume using 10ul fresh beads + 50uL PEG-NaCl buffer (20% PEG + 2.5 M NaCl), eluted and quantified by nanodrop. The final PCR was conducted with 0.1ng of this sonicated and purified RCA product as input, for 8 cycles at 55°C annealing followed by 6 cycles at 72°C annealing. This template amount was carefully optimized to avoid barcode switching. The completed reaction was run slowly on a 2% TAE gel and progress was monitored until the 90bp “barcode only” species ran off the gel to facilitate gel purification of the 362bp “barcode-intron-barcode” species using Zymo Gel DNA Purification Kit.

### Assembly of the final plasmid library

The barcode-intron-barcode product was mixed at a 5:1 ratio with the GFP fragment of the library acceptor plasmid and subjected to 15 cycles of Golden Gate Assembly (GGA) using the enzyme BspMI. The resulting product, referred to as the Library Vector (LV), was SPRI-purified, diluted to 50ng/uL for electroporation into 8 aliquots of Endura electrocompetent cells, using 2µL DNA per 25µL cells. These cells were allowed to recover and then plated at various dilutions on 8×15cm plates with Ampicillin (100µg/mL) and incubated at 30°C overnight. The next day all plates were scraped and DNA was purified via Zymo DNA MaxiPrep Kit. The purified LV was digested with PsrI and gel purified to linearize it between the intron and the first of the two barcodes, at the site previously introduced upstream of the intron 5ʹSS. The pre-digested dTomato fragment was added at a 2.5:1 ratio and the two fragments were joined with 15 cycles of PsrI GGA assembly using PsrI. This final product, referred to as RIL for Random Intron Library, was SPRI-purified and electroporated as previously described for LV, this time plated on agar with Ampicillin (100µg/mL) and Cotrimoxazole (20µg/mL). Plates were incubated at 30°C overnight and colonies were scraped and purified as described before. This RIL was used for an initial round of experiments (RNA-seq replicates 1-5), after which the same plasmid library was re-transformed at several different dilutions to bottleneck the diversity of intronic variants down to ∼50,000. This iteration of the library was transformed into the methylation-deficient C2925 *E. coli* strain following manufacturer’s protocol and the transformation yielding ∼50,000 colonies (as determined by counting colonies on further dilution plates) was selected to be scraped and maxiprepped as described above.

### Generation of pooled cellular library

For the first round of experiments (replicates 1-5), 75µg of the initial Random Intron Library was transfected into 43 million HEK293T A2 cells in nine 12-well plates using Lipofectamine 2000, following the protocol described in (Khandelia et al., 2011). For the second round (replicates 6-10), 320µg of the bottlenecked RIL plasmid library was pre-mixed with1.6µg Cre recombinase vector and 40ng of each spike-in control plasmid (i.e. intronless barcodes, UbC intron). The integration was further scaled-up to transfect 80µg of pre-mixed plasmids with 80µL Lipofectamine 2000 into each of four 15cm plates with ∼20 million HEK293T A2 cells each. The puromycin selection was performed as before. Approximately half of the resulting pool of cells was immediately frozen into aliquots with ≥5 million cells/vial and the other half cultured, with at least 5 million cells per split to avoid bottlenecking the library. The total RNA was extracted from the pooled cells using Zymo Quick RNA Miniprep kits in five replicates for downstream processing.

### Long-read sequencing and analysis

#### Sample preparation and sequencing run

Two preparations of the bottlenecked RIL plasmid library were linearized in four digestion reactions, each with one of four restriction enzymes (EcoRI, SalI, XbaI and PsiI), which were subsequently SPRI purified and pooled for sequencing on one Oxford Nanopore R10 PromethION flowcell at the MIT BioMicroCenter.

#### Barcode-to-intron dictionary generation

Minimap2 was used to align the Nanopore reads to the plasmid backbone and extracted the intron and barcode sequences from all reads mapping contiguously between the dTomato and GFP barcodes. With a relatively high (∼4% in expected constant regions) error rate in long-read sequencing, each 160nt intron random region is expected to have 6 mutations on average, so we rely on seeing each intron multiple times to confirm its sequence. We clustered the intron sequences with the UCLUST algorithm (Edgar, 2010) and stored the consensus intron sequence and most common barcode in a provisional barcode-int dictionary, discarding clusters with fewer than 3 reads. In principle, we allowed introns to have multiple barcodes, but in practice different barcodes mapping to the same intron were always closely related in sequence, and thus likely resulted from sequencing errors in reading of the same barcode sequence. We later validated the accuracy of this barcode-intron map by confirming that the intron sequences in unspliced RNA seq reads were as expected for that read’s barcode.

#### Barcode pairing validation

We validated the integrity of barcode matching between dTomato and GFP in the library by considering each aligned read from the Nanopore sequencing and computing the Levenshtein distance between the two barcodes. We tallied the rate of matches (Lev dist ≤4) and mismatches for each barcode and kept only those that have 10% or fewer mismatches. All other barcodes were discarded from the dictionary and not considered in further analyses.

### RNA sequencing and analysis

#### Sample preparation and sequencing run

For each RNA-seq sample, 2µg RNA extracted from the pooled random intron library was treated with Turbo DNase (Thermo) to remove any residual genomic DNA before further processing. We removed DNase using Zymo RNA clean & concentrate kit and next performed gene-specific RT using SuperScript IV (Thermo) and two primers, complementary to the GFP and dTomato N-termini, respectively, in order to generate cDNA of the two reporter gene 5ʹUTRs only. After first strand synthesis we treated the samples with RNase cocktail (Thermo) to avoid interference with downstream PCR. The cDNA was amplified with 16 cycles of PCR using Q5 polymerase (NEB) and custom sample-specific indexing primers. The PCR product was purified by a double-sided SPRI purification, first removing genomic DNA with 0.55x volumes of beads and then adding 85µL beads for a final 1.4x volume. The amplicon was sequenced with either 50+50nt PE reads for replicates 1-5, on an Illumina HiSeq at the Whitehead institute sequencing core, or 105+45nt PE reads for replicates 6-10 using the Element AVITI instrument at the MIT BioMicroCenter.

#### Read processing

Reads were first scanned for a correct sample index and for a match or close match to the constant sequence distinguishing the GFP from dTomato 5ʹUTR, and discarded if they were missing either of these. GFP reads were next classified into spliced, unspliced, or other by searching for a match to the expected exon-exon junction sequence. Unspliced reads with the expected intron sequence (constant 5ʹSS and 3ʹSS) were counted for that barcode as unspliced, while reads matching neither the expected spliced sequence nor the expected unspliced sequence were set aside for further analysis. Finally, classified reads were filtered for perfect or close matches to known and trusted barcodes and otherwise discarded. This yielded ∼125 million RNA-seq reads across 10 biological replicates.

#### Splicing and cryptic splicing analysis

Reads identified as originating from GFP were classified into spliced or unspliced based on the presence or absence of the expected exon 2 sequence TCGTGAA 3 nt downstream of the exon-exon junction. Unspliced reads were further categorized into canonical unspliced if they instead had the predicted sequence of the unspliced intron (AGTAGCG), or unknown isoform if they had a significantly different sequence. The splicing efficiency of each intron was obtained by tallying the fraction of correctly spliced reads per barcode, per sample, taking the median across all samples with at least 100 GFP reads. To identify instances of cryptic splicing, reads with unusual intronic sequence were partitioned into those with mismatches at the 5’end or 3’end (some reads belonging to both sets) and a window of 16 or 20 nt (for 5’ or 3’ respectively) from the read was used to scan along the predicted intron sequence, with Levenshtein distance calculated at each position. If all reads for a given barcode had an optimal match at the same position of the intron, and the Levenshtein distance between the read and intron at that position was ≤2, this position was identified as a cryptic splice site.

#### Iterative FACS analysis

Fluorescence-assisted cell sorting (FACS) experiments were performed at the Koch Institute Flow Cytometry core using the BD FACSAria III instrument (BD Biosciences). Cells were prepared and sorted as described in Flow Cytometry methods section. For sort 1, 60 million pooled integrated random intron library cells were sorted in 2 replicate batches. Bins were drawn to select the top and bottom 10% of cells along the diagonal FITC versus PE signal axis and 1.8–2.6 million cells were collected per bin per replicate, into tubes containing 50% DMEM and 50% FBS. These cells were plated in 4×15cm dishes and allowed to grow for 4 days. After trypsinizing each plate from the first sort, cells were counted and 1 million cells from each plate were pelleted and snap-frozen in an aluminium block cooled to -80°C for later RNA extraction. Bins were re-drawn to capture the top 10% of the green-shifted populations and the bottom 10% of the red-shifted populations, and 0.6–0.7 million cells were collected per bin per replicate. These were plated in T25 flasks and allowed to grow for an additional 3 days before the final sort, at which point 0.15 million cells from each flask were pelleted and snap-frozen. The remainder of each flask was sorted into bins were re-drawn to enrich for the most extreme 10% once again, yielding 130–180 thousand cells per bin per replicate. These were immediately pelleted and snap-frozen such that downstream processing of all 12 samples from the iterative FACS experiment occurred in parallel.

#### IME analyses

For bulk RNA-seq we filtered all barcodes to consider only those detected in three or more samples, then ran DESeq2 to get intron-specific log2fold change estimates, using Ashr shrinkage (Love et al., 2014; Stephens, 2017). For the iterative FACS libraries, we identified bins of introns detected in both trajectories, and designated “green” and “red” sets as those present in every stage of green sorting and no stage of red sorting, and vice versa. These sets in combination were used with mRNA-level IME scores to study the splicing and sequence features of introns as a function of their magnitude of gene expression enhancement.

#### Random introns +/– polyU mini-library

The second library was screened using methods similar to the random library with a number of modifications. Details of intron selection and cloning workflow available upon request. Procedural differences are as follows:

- 12nt barcodes composed of all 4 nucleotides were iteratively randomly generated such that none contained an ATG, a restriction site used in the library assembly, or a mononucleotide run longer than 4, and each new barcode added to the list had Levenshtein distance ≥3 from all previous barcodes.
- Removed/mutated the potential 5ʹSS present in the 3ʹ-end of GFP and dTomato
- Used pMBYS124, a plasmid modified from pEK1, to insert the introns and re-cloned the dTomato fragment into Kanamycin-resistant plasmid to enable antibiotic selection at the 2nd step of library cloning
- RNA was extracted after transient transfection with no Cre or puromycin selection

#### Transient transfection

8×10^6^ HEK293T A2 cells or HeLa A12 cells were plated in 30mL media on 15cm dishes, in two replicates each on the day before transfection. The following day each plate was transfected with 40µg second intron library plasmid DNA, using Lipofectamine 3000 by following manufacturer’s instructions. After 24h of transfection, the media was removed and cells were rinsed twice with PBS, trypsinized and split 1:3 into 2 new 15cm dishes. After another 24h (48h of transfection total) these cells were trypsinized and total RNA extracted using TRI-reagent and Direct-zol RNA Miniprep Plus kit.

#### RNA-sequencing library prep & analysis

Extracted total RNA was converted to cDNA with SuperScript IV (Life Technologies) using primers specific to the GFP and dTomato coding regions so as to amplify only the two reporter gene 5ʹUTRs. These cDNA samples were amplified by 12 and 15 cycles of PCR for HEK and HeLa cells respectively. Samples were dual indexed with custom primers and sequenced on the Element AVITI instrument at the MIT BioMicroCenter. RNA-seq reads were classified first into samples and then into one of three types (dTomato, spliced GFP, unspliced GFP) based on matching short sequences at expected locations in the sequenced amplicon. Barcodes were extracted and only reads with barcodes matched to introns from the designed library were kept. For subsequent analyses barcodes were filtered to include only those represented by at least 100 reads per replicate to remove noise from low count data.

#### Simulation

We modelled variation in GFP and dTomato read counts across replicates assuming different ranges of underlying read ratios, i.e., IME strengths. We sampled read counts for each intron from the empirical distribution of read counts for introns in our data, and split them into GFP/dTomato by sampling from a binomial distribution with probability determined by that intron’s expected read ratio. Underlying transfection-specific noise was injected by sampling dTomato read count only once per transfection per intron and using the same values for both replicates. We included parameters “noisiness” and “txf_noisiness” to describe additional transfection-independent and -dependent noise, which were each computed as addition of an integer drawn from a normal distribution with spread equal to the initial read count times the noisiness parameter (ultimately 0.25). Finally, we added a “txf_noise_sparsity” parameter controlling what fraction of introns had transfection-specific noise applied to them. These parameters were manually adjusted until the resulting scatter plots resembled the real data as closely as possible.

### Data Availability

The RNA-seq data underlying this article are available in the NCBI GEO Database, under accession code GSE278584. The primary analysis code is available at https://github.com/ejkk0/IME, with other scripts available upon request.

## Results

### Different introns enhance gene expression to different extents

We first sought to reproduce the observation from plants that different well-spliced introns can stimulate different levels of expression from otherwise identical reporter genes. We used the HILO-RMCE system (Khandelia et al., 2011) for rapid generation of human cell lines with single-copy genomic integration of our reporter constructs. We selected five introns which had been reported in previous literature to enhance gene expression (Fig. 1B) and cloned each separately into the same position in the 5ʹUTR of an enhanced green fluorescent protein (EGFP) gene driven by the human UbC promoter. Each of these vectors, plus an otherwise identical intronless EGFP vector, was co-transfected with a Cre recombinase plasmid into HILO-RMCE HEK293T A2 cells (Fig. 1C), which were then maintained under puromycin selection for two weeks to ensure integration of the transfected DNA.

Each intron-containing cell line yielded higher steady state levels of EGFP mRNA and protein than the intronless version, as measured by qRT-PCR and flow cytometry (Fig. 1C,D), and RT-PCR analysis of EGFP RNA from these transgenic cell lines showed that each intron was efficiently spliced (Fig. 1E). The magnitude of IME – defined as the fold increase in expression over intronless – varied from about 4-fold to more than 10-fold at both mRNA and protein levels. Interestingly, across introns, the increase in protein yield was highly correlated to the increase in mRNA. The variability in IME attributable to these introns suggests that intron identity (i.e. sequence) contributes to differences in enhancement. For the purposes of this study, we defined IME as the increase in expression of a host gene attributable to the splicing of an intron. Thus, if an intron were to enhance expression by containing a transcriptional enhancer, or by stabilizing unspliced transcripts, for example, this would not be considered IME.

### Design and construction of a barcoded dual-reporter system to study IME

In order to investigate the relationship between intron sequence and IME, we conceived a large-scale reporter experiment to measure the enhancing activity of many introns in parallel (Fig. 2A). We reasoned that sequence-specific differences in IME may be driven by the recruitment of sequence-specific RNA-binding proteins (RBPs), which typically bind short, 4-8 nucleotide (nt) motifs within introns. Using a library of comparatively long (160 nt) random sequences allows exploration of the sequence-activity relationship of IME in an unbiased manner, with a large diversity of intron sequences and coverage of all short motifs in different relative arrangements.

We generated a DNA library of introns with 160 nt of fully random internal sequence, flanked by strong splicing signals (UbC 5ʹSS and EF1a branch point and 3ʹSS) for an intron length of 212 nt in total. We inserted these introns into the UbC promoter and 5ʹUTR in place of the endogenous UbC first intron, upstream of the EGFP coding sequence. Each intron was randomly paired with a random H_18_ barcode (18 nt comprising A, C or T, but no G) located in the 5ʹUTR downstream of the intron. Locating the barcode in a noncoding exonic region of the transcript enables recovery of the intron sequence from RNA sequencing reads of the spliced mRNA, and the exclusion of Gs precludes the formation of alternative splice sites as well as upstream AUG start codons, which are known to be a major 5ʹUTR sequence determinant of translational efficiency and mRNA stability (Hurt et al., 2013; Sample et al., 2019).

A key feature of our design is that we included a second minigene in the reporter system, the red fluorescent protein dTomato (dTom), roughly 1 kb upstream of the transcription start site (TSS) of EGFP (Fig. 2A). We constructed the library so that the dTom and EGFP 5’UTRs in each plasmid bear identical H_18_ barcodes, and therefore the ratio of read counts originating from EGFP versus dTomato for a particular barcode serves as a measure of the associated intron’s impact on EGFP expression. This design controls for: (i) the abundance of the plasmid in the library; (ii) the transcriptional activity of the reporter cassette in the particular cell(s) that integrated it; and (iii) any impact of the barcode on mRNA or protein expression from the transcript.

Accurate duplication of randomly generated barcodes in a pooled fashion had not been accomplished previously to our knowledge. We ultimately achieved this through circular ligation of the intron-barcode pairs, followed by rolling circle amplification, fragmentation, PCR, gel electrophoresis and subsequent size selection for fragments containing two copies of each barcode flanking a single intron (Fig. 2B; Methods). We optimized this procedure to reduce barcode mismatching from PCR-induced chimerism (Hegde et al., 2018) and generated a highly diverse vector library containing over 300,000 unique introns detectable by short-read sequencing of plasmid DNA. We proceeded to investigate IME using both this full library as well as a bottlenecked (sub-sampled) library ∼10% of the original size, in order to increase the read depth associated with each individual intron.

We assayed the reduced plasmid library via Oxford Nanopore long-read sequencing to construct the barcode-to-intron map and to validate the integrity of dTom-GFP barcode matching. For each intron, we analyzed the set of associated barcodes and excluded promiscuous barcodes which could not be confidently assigned to a single intron (Fig. 2C). This procedure yielded a dictionary of 49,737 barcode-intron pairs, whose intron sequences were supported by clusters of 116 reads on average (median 78). We additionally confirmed whether, in each read, the dTom barcode matched the GFP barcode as intended. We expected some degree of mis-pairing to have occurred in the library cloning procedure and sought to restrict our analysis to only those barcodes that were correctly paired. We found that 38,158 GFP barcodes (76.7%) were paired with matching dTom barcode in ≥90% of plasmid reads (Fig. 2D). All further analyses focused on this set of confident barcodes and introns only.

### Short random introns are efficiently spliced in human cells

We co-transfected this plasmid library with a Cre recombinase vector into the HILO-RMCE HEK293T A2 cell line (Khandelia et al., 2011) in order to create a pooled cellular library with single-copy genomic integration of the random-intron-containing EGFP. As a readout of intron effects on steady-state RNA level, we extracted total RNA from these cells and sequenced an RT-PCR amplicon of the GFP and dTom 5ʹUTRs. In each sample, this experiment yields three types of reads per barcode: dTom reads, spliced GFP reads, and unspliced GFP reads (Fig. 3A). In total we performed ten biological replicates of RNA extraction from the cellular library for amplicon sequencing, five with the full plasmid library and five with the bottlenecked version. It was essential to determine whether random introns would be efficiently spliced in these cells, despite their internal sequence being random and thus lacking natural splicing regulatory elements other than the core 5’SS, branch point and polypyrimidine tract/3’SS. Across the ten replicates of RNA-seq, over 98% of GFP reads were spliced as expected, using the constant 5ʹSS and 3ʹSS present in all introns (Fig. 3B).

**Figure 3.**
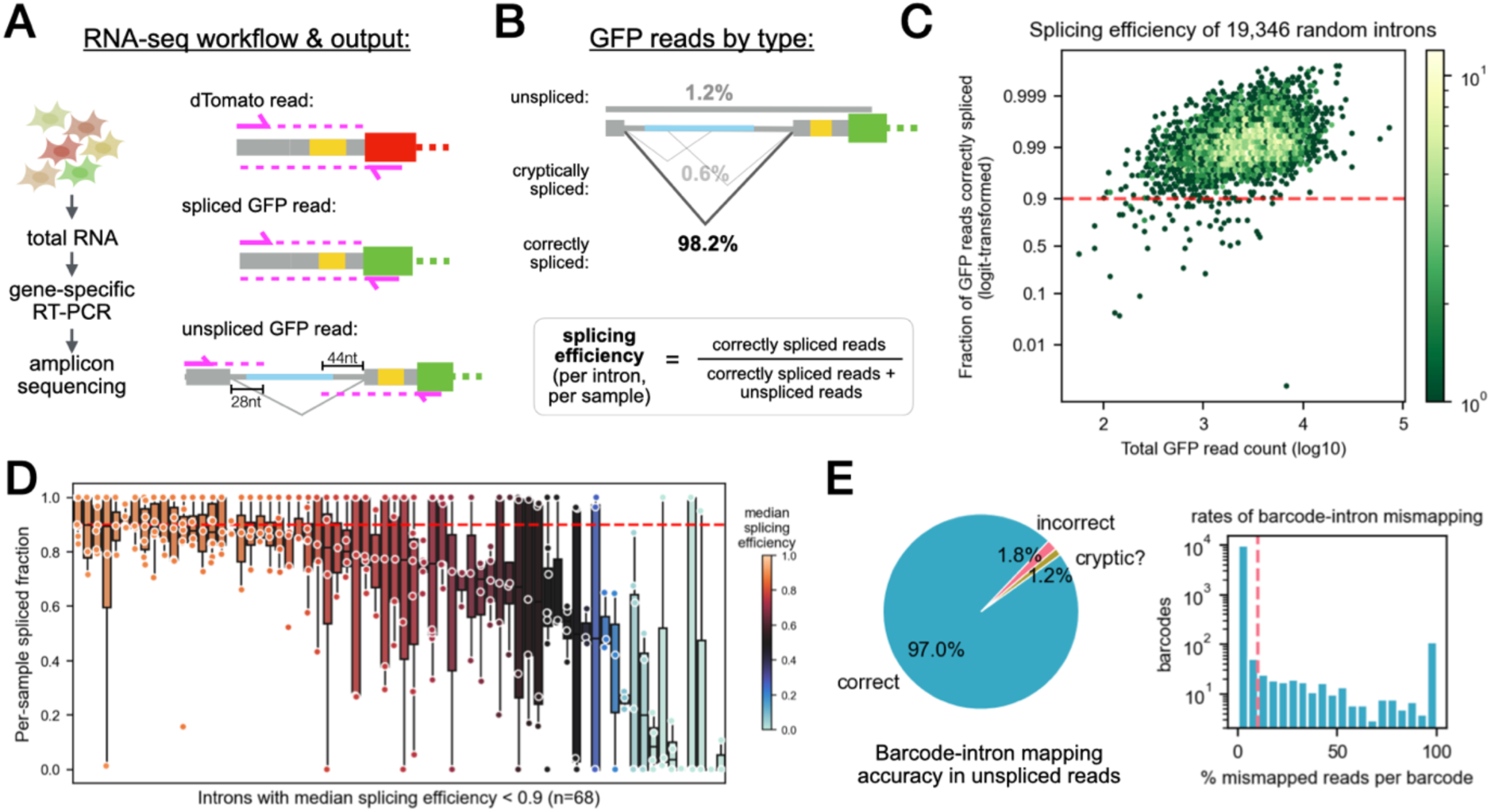
Almost all short random introns splice efficiently. **A)** Schematic of RNA-seq workflow and resulting read types. **B)** Of all the GFP reads detected, the vast majority are spliced as expected between the intended library splice sites. Splicing efficiency calculations per intron, per sample take into account correctly spliced reads and unspliced reads that look as expected, disregarding reads of unknown splicing status. **C)** Median splicing efficiencies of all introns in the random library, across all samples in which that intron has at least 10 GFP reads, as a function of total GFP read count for that intron. **D)** Per-sample splicing efficiencies of introns with median splicing efficiency <0.9. Boxplot centers represent medians, box edges represent interquartile range (IQR) or middle 50% of data, whiskers extend to 1.5x IQR past each box edge. **E)** Fraction of barcodes for which intron random regions in unspliced reads match the expected sequence. To be classified as “correct,” the intron must match in ≥90% of reads with the associated barcode.

Unspliced reads were associated with 10,507 introns, but usually accounted for a small fraction of the GFP reads from any individual intron. Of 19,346 introns that met our minimum read count threshold for analysis, requiring at least 10 GFP and 10 dTom reads in at least three samples, the vast majority (99.7%) were well-spliced, with median splicing efficiency 0.9 or higher across all samples with minimum 10 GFP reads (Fig. 3C). Only 68 introns had median splicing efficiency below 0.9, and the majority of these were still above 0.8 (Fig. 3D). Because the PCR step in our amplicon-sequencing approach likely favors shorter (spliced) products over unspliced, we do not report percent spliced in (PSI) values here; however, when directly compared to a natural intron control which is known to be well-spliced (UbC), the random introns exhibited similar observed splicing efficiencies (Fig. S2F). These observations suggest that presence of strong splice sites and a branch point, as in our reporter, may be sufficient for splicing of short (∼200 nt) introns, more or less independently of the intervening sequence (De Conti et al., 2013).

The unspliced reads also serve a valuable quality control role for this system: in the latter five of the ten RNA-seq replicates performed, our sequencing scheme read 28 nt past the fixed 5ʹSS and 44 nt upstream of the fixed 3ʹSS in each transcript (Fig. 3A). Accounting for constant regions, this yields 16 nt and 4 nt, respectively, from the random region of the intron on either end, allowing us to verify that the intron sequence is that expected from the barcode in the read. In a first pass analysis, just over 97% of barcodes observed in unspliced reads (10,194 out of 10,507) correctly predicted the first 16 nt of intron sequence with ≥90% accuracy (Fig. 3E), confirming high accuracy of barcode-intron mapping in our system. Of the remaining 3% of “mismapped” reads, we noticed that most appeared to be truly unspliced and mismapped, while a subset aligned partially to the correct intron sequence and appear to be spliced to a cryptic 5ʹSS or 3’SS created by chance in the random region of the intron. These reads were therefore reclassified as cryptically spliced rather than mismapped, increasing our estimate of the accuracy of barcode-intron mapping to almost 98%.

### Usage of cryptic 5ʹ and 3ʹSSs

To explore the distribution of cryptic splicing in the random library, we investigated the set of ∼250K GFP reads that were neither unspliced nor canonically spliced. The constant 5’ and 3’SS are relatively strong, with MaxEnt scores (Yeo and Burge, 2004) of 10.7 and 13.0 bits, respectively (Fig. 4A). Assuming cryptic splicing from one of these SS to an internal cryptic site, we reasoned that we should be able to infer the exact location of the cryptic site from the read sequence. Splicing to an internal cryptic 5’SS would generate reads that included exon 1 and a portion of the 5’ end of the intron, followed by exon 2, and vice versa for splicing to an internal cryptic 3’SS (Fig. 4B). Reads with unexpected sequence at both ends were largely derived from cryptic sites in the intron constant regions, i.e. close enough to each end to be captured in both reads (Fig. S1A). Altogether, about 1 in 4 unspliced reads had some unexpected sequence at either or both ends (Fig. 4C).

**Figure 4.**
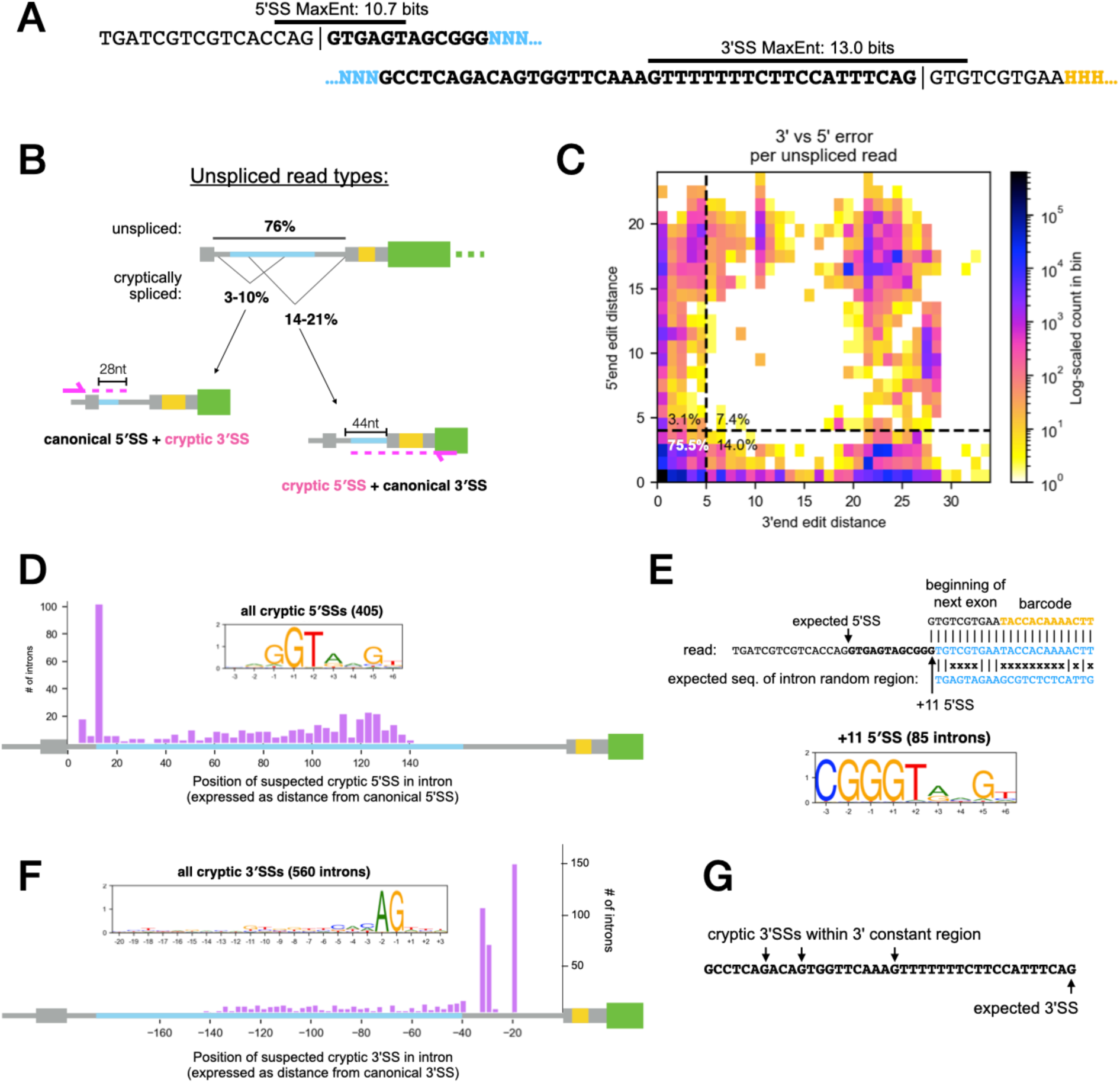
Cryptic splicing in random introns. **A)** Splice sites used in random library, annotated with respective MaxEnt scores. Intron-exon boundaries denoted by |, constant 5ʹ and 3ʹ SS sequences flanking random region in bold, intron random nucleotides in blue, barcode random nucleotides in yellow. **B)** Schematic of unspliced GFP read types, i.e. completely unspliced vs. 5ʹ or 3ʹ end cryptic splicing. **C)** 2D histogram of edit distance from expected sequence at 5ʹend and 3ʹend of all unspliced reads. Percentage of reads in each quadrant are annotated at inside corners. **D)** Distribution of locations within intron where evidence is seen for use of cryptic 5ʹSSs. Inset: sequence logo of nucleotides [-3,+9] at these positions. **E)** Example of an unspliced read inferred to have used the +11G as a cryptic 5ʹSS, and sequence logo for the subset of 5ʹ cryptic splicing events mapping to this position. **F)** As in D, for cryptic 3ʹSS. **G)** Constant sequence at 3ʹend of random library with positions of suspected cryptic 3ʹ splicing indicated. See also Supplementary Figure S1.

By aligning the 3ʹ end of each of these reads with the expected intron sequence, we were able to identify over 400 likely cryptic 5ʹSS (Fig. 4D, S1B). Overall, aside from one exceptional position, the frequency of cryptic 5’SS increased with distance from the canonical 5ʹSS up to 150 nt, and the inferred 5’SS motifs resembled the canonical human 5ʹSS motif, as expected (Fig. 4D). The exception was the proximal +11G position in the 12 nt of 5’ constant sequence, which was a hotspot of cryptic splicing, with 85 introns using the +11G in at least 10 reads. This site is likely preferred because it already has consensus bases at the –3, –1 and +1 positions of the 5’SS motif for a 5’SS at +11 in the intron (Fig. 4E). The absence of cryptic 5’SS past 150 nt implies a minimum intron length of ∼70 nt, consistent with previous studies in mammals (Wieringa et al., 1984).

Repeating this procedure with reads that had matching to the 3ʹ end of their intron, we identified instances of introns using internal cryptic 3ʹSS (Fig. 4F, S1C). These, too, were distributed throughout the intron no further than ∼150 nt upstream of the canonical 3ʹSS, again implying a minimum intron length of ∼70 nt. These sites were enriched for positions within the constant region that already matched the minimal 3’SS consensus NAG (Fig. 4F,G). Curiously, we observed a negative correlation between the scores of cryptic splice sites and their positions within the random region of the intron, with weaker cryptic 3’SS and 5’SS motifs being observed only near the 3ʹ end of the intron (Fig. S1D,E), even when splice sites overlapping constant regions were excluded. This suggests that the constant regions at the 3’ end of the intron and the 5’ end of exon 2 may represent a particularly strong context for splicing.

### Random introns enhance gene expression at mRNA and protein levels

Having confirmed that the introns in our library predominantly splice as expected, we next examined the GFP and dTom read counts per barcode to study each intron’s effect on GFP mRNA expression. For this analysis we used all ten replicates of amplicon sequencing, across which the mean GFP and dTom read counts of the different barcodes varied over 5 orders of magnitude (Fig. 5A). Negative (intronless) and positive (UbC intron) controls were spiked into the random intron plasmid library at a 1:5000 mass ratio prior to transfection and, as expected, yielded total read counts roughly ten-fold higher than the library average. Batch variation in the global average ratio of GFP to dTom reads was observed, corresponding to the two sets of replicates performed pre- and post-bottleneck (Fig. S2A). To control for these batch effects and remove noisier (lower-count) barcodes, we applied a minimum read count filter and normalized read counts across replicates using DESeq2 (Love et al., 2014). We reasoned that the question of intron effects on GFP expression relative to dTom can be treated as a differential expression analysis, where GFP and dTomato counts from the same replicate are handled as paired “treatment” and “control” samples, and we model the variation across replicates to determine which “genes” (distinct introns in our analysis) are significantly altered in the treatment versus control.

**Figure 5.**
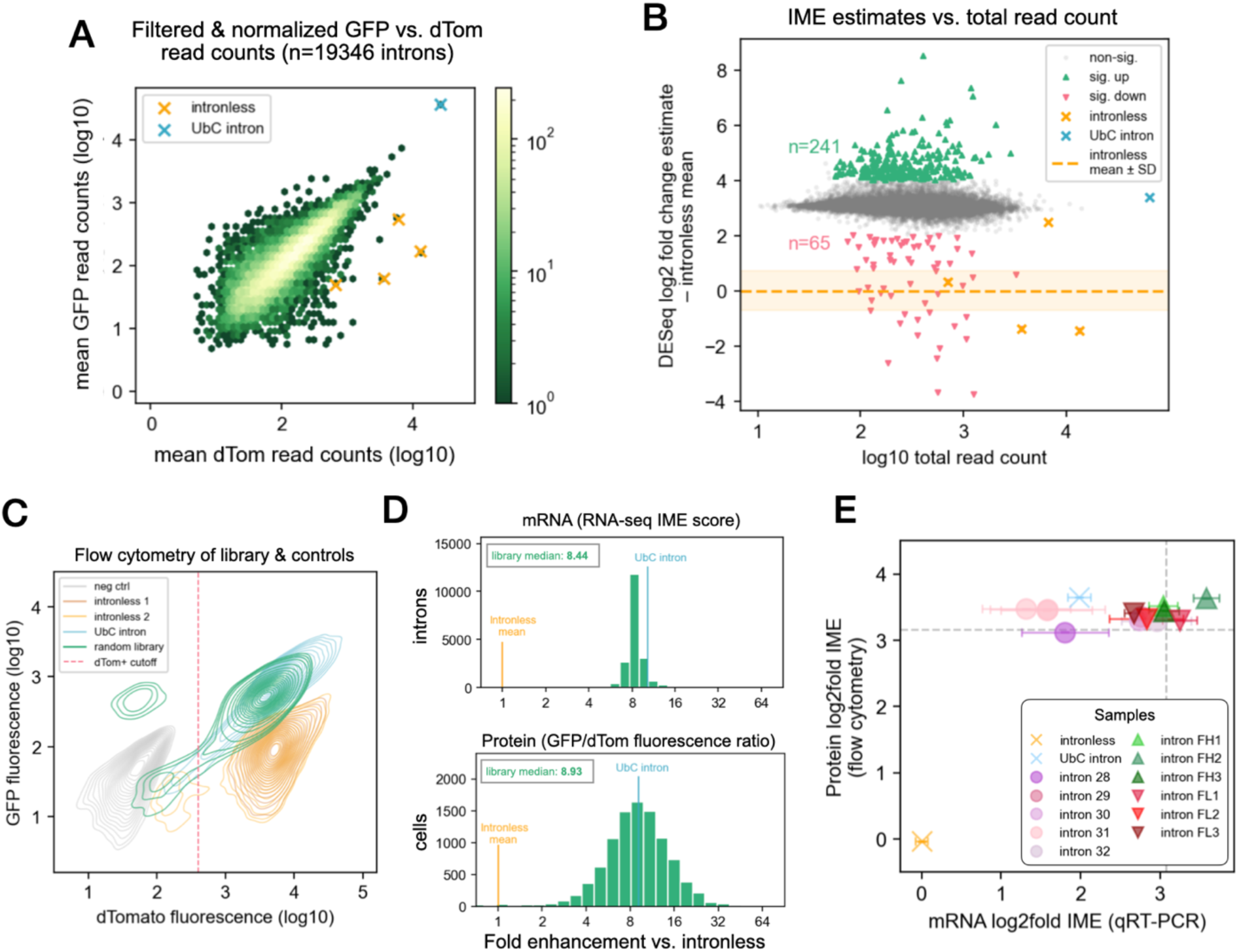
Random introns enhance GFP mRNA and protein expression. **A)** GFP and dTomato RNA-seq read counts per intron, mean across ten samples. **B)** DESeq2 log2 fold change estimates vs. total read count for each intron. Estimates are normalized by subtracting the mean log2 fold change of the four intronless control barcodes. Introns called significantly different from the mean of the library are highlighted (*p≤*0.05, absolute log2 fold change ≥1). **C)** Flow cytometry of cells with integrated library overlayed with various controls. For calculation in D, library cells not expressing dTomato (red dotted line) were excluded. **D)** Fold enhancement at mRNA and protein level relative to intronless are compared on the same scale. Steady-state GFP mRNA and protein levels are each ∼eight-fold higher with a short random intron than with no intron. **E)** qRT-PCR and flow cytometry of transgenic cell lines expressing GFP with individual selected introns for validation. Error bars denote standard error of the mean. See also supplementary Figure S2.

The filtered and normalized read count data suggested that the presence of an intron alone, independently of its sequence, stimulates GFP mRNA expression (Fig. 5B). Since the GFP and dTomato counts are treated as separate libraries during DESeq2 modeling and shrinkage, their mean ratio across all barcodes is set to zero during normalization. To estimate the magnitude of each intron’s effect on expression compared to an intronless version of the same gene, we subtracted the mean log ratio of the four intronless controls, yielding a library average log_2_ fold change of ∼3.1, or an absolute fold increase in GFP mRNA of ∼8.5-fold compared to intronless GFP. To validate this observation, we also measured the difference in GFP/dTom ratio by qRT-PCR of RNA extracted from a pool of cells with the whole library integrated compared to cells with integration of the intronless plasmid, yielding an estimated average enhancement of 5-fold (Fig. S2B). These two approaches for measuring the aggregate IME of the library have distinct strengths and weaknesses, as discussed below. Nearly all intron-containing reporters exhibited a higher GFP/dTom ratio than the four intronless reporters, indicating that considerable enhancement of expression is a general property of splicing in this context.

Some variation in enhancement levels occurred within the library: 244 out of 19,346 introns analyzed had significantly stronger IME than other introns, while 65 were had significantly weaker IME, including 3 of the 4 intronless control barcodes (Fig. 5B). Prior to application of DESeq2, we observed that GFP/dTom read count ratios were not always well correlated between replicates: specifically, replicates from separate transfections conducted in parallel had minimal correlation, while replicates originating from the same transfection were highly correlated (Fig. S2C). Nevertheless, the intronless control barcodes consistently yielded significantly lower GFP/dTom read count ratios than the rest of the library, across all replicates, whether between or within transfections. This observation implies that our setup can reveal reproducible variation in IME as long as it is of sufficient magnitude.

To estimate the range of IME values represented by the data, we performed various simulations of our library experiments. These simulations modeled both the sampling noise in estimation of individual IME values from RNA-seq and the variability resulting from different transfections. This includes any experimental variability in the precise transfection and subsequent antibiotic selection conditions, as well as epigenetic/cell state differences between cells that integrate a particular intron across different transfections (Supplemental Note 1). A simulation which visually recapitulated the spread and density of points in the observed scatter plots was obtained (Fig. S2D). With this simulation, the correlations observed between replicates matched best to simulated values with a total relative IME score log ratio range of 0.1-0.3 (Fig. S2E). This is equivalent to ∼95% of introns in the library conferring log_2_ enhancement between 2.7 and 3.5 (i.e. IME between 6.5-fold and 11-fold). Taken together, these data suggest that virtually any well-spliced intron can confer strong IME.

To assess whether this effect also manifested at the level of protein expression, we used flow cytometry to measure the GFP and dTomato fluorescence in the pooled cellular library, alongside our UbC intron and intronless control lines, and negative control cells expressing neither protein (Fig. 5C). The random library and the UbC intron control both displayed similar dTom expression and significantly higher GFP expression than the intronless control lines. We observed a small population of dTom-negative cells in the library and verified that these represent carryover of a cloning intermediate; these barcodes are excluded from subsequent analyses via removal from the barcode-to-intron dictionary, and from this analysis by considering only dTom+ cells (Fig. 5C, red dotted line) in calculating the average expression ratio of the library.

We computed the per-cell ratio of GFP to dTom fluorescence after subtracting autofluorescence (median intensity of negative control cells) and compared the distribution of ratios in the library to that of ratios in the intronless control cell lines. Consistent with the RNA-seq-derived estimates, we found that cells with intron-containing GFP produce on average 8.9-fold more GFP than intronless cells when each is normalized to its paired dTom measurement (Fig. 5D). In both cases, the UbC intron falls close to or just above the mean of the random introns. Though the distribution of mRNA-level effects appears highly concentrated compared to the protein-level measurements, this difference in spread likely reflects differences in measurement noise and data processing (e.g., the shrinkage and normalization performed at the mRNA level) rather than biology. Furthermore, the flow cytometry experiment does not assess which intron is in which cell, only capturing the bulk population, so the two distributions are not directly comparable.

We sought to validate these high-throughput measurements by cloning and testing the IME activity of individual random introns from the library. We selected 11 introns and constructed 11 transgenic lines, each harboring GFP containing one of these introns and an intronless dTom, without barcodes. We assayed their mRNA- and protein-level enhancement by qRT-PCR and flow cytometry and compared it to the expected values from the high-throughput measurements as well as the intronless and UbC control lines (Fig. 5E). We selected introns with a range of predicted IME strengths, and observed a range of mRNA-level IME values from 2.5- to 12-fold, similar to the ranges predicted by DESeq2 and by our simulations. For a subset of these introns, efficient splicing was confirmed using RT-PCR (Fig. S2F). For one specific intron, we conducted three separate transfections to produce three independent cell lines, verifying that the measured IME value is reproducible (Intron 31, Fig. 5E). Discrepancies in the relative ranking of introns measured by qRT-PCR versus RNA-seq suggest the presence of biological variability between the pooled and individual transgenic lines.

### Selection of introns with stronger and weaker IME by iterative FACS

In order to identify the specific random introns with stronger and weaker protein-level IME, we devised an iterative fluorescence-assisted cell sorting (FACS) procedure to enrich for introns with extreme enhancement values. We collected cells from the library with the highest and lowest GFP/dTom fluorescence ratios by empirically slicing the top and bottom 10% of cells along the *y=x* diagonal (Fig. 6A, S3A). After each sort, we reserved some sorted cells for RNA extraction and sequencing, and cultured the remainder until re-sorting several days later. Three successive sorts in two independent trajectories led to smaller and smaller sets of barcodes being captured, with fluorescence intensity distributions shifted higher or lower, as expected (Fig. S3B,C).

**Figure 6.**
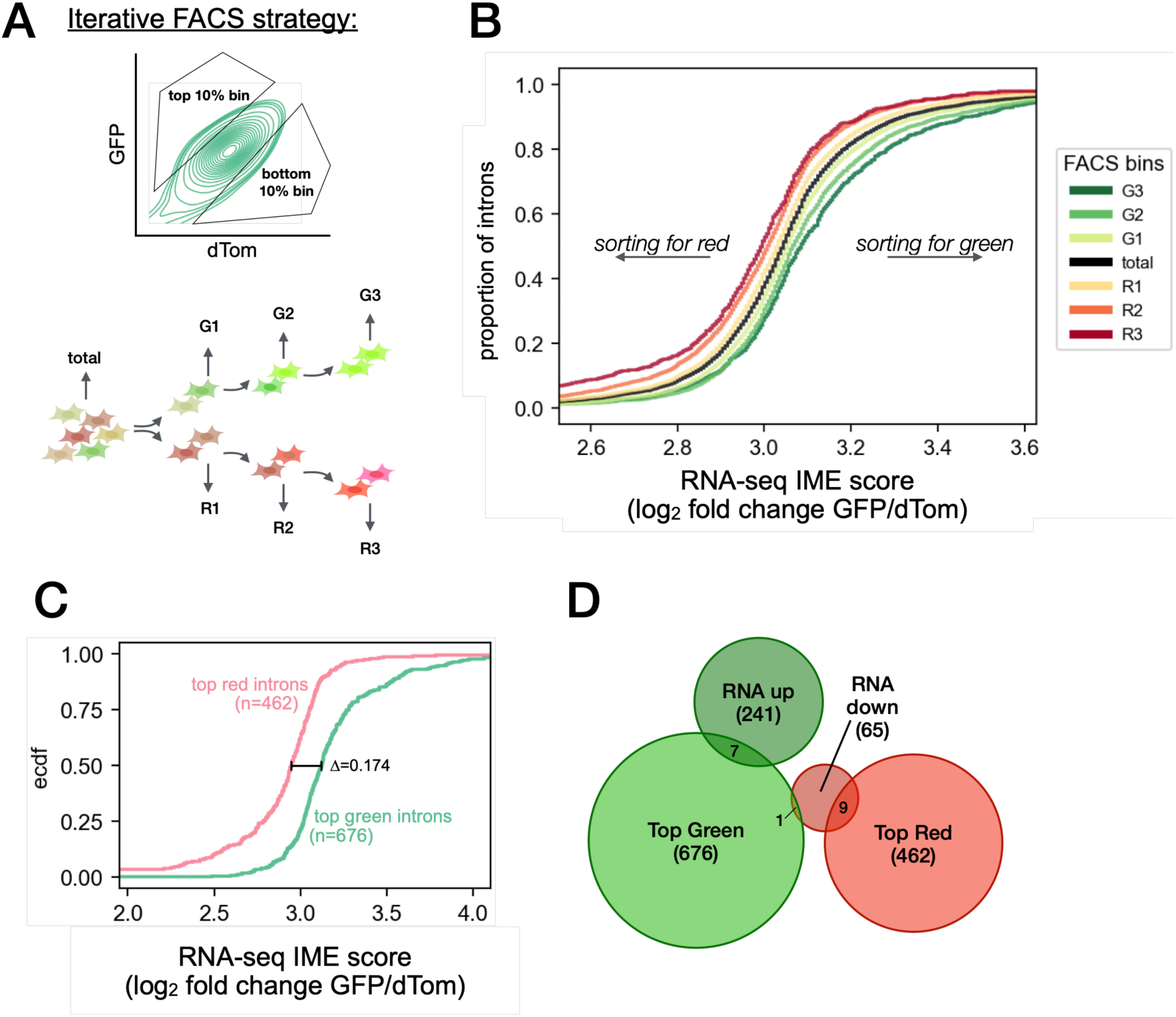
Iterative enrichment of cells with extreme GFP/dTomato fluorescence ratios allows study of protein-level IME. **A)** Schematic of iterative FACS experimental design. **B)** CDFs of mRNA-level IME scores (DESeq l2fc estimates) for the set of introns detected at each sort stage. Shown are the union of both replicate trajectories, using 100 reads as cutoff for detection. **C)** Designated sets of introns with strongest and weakest protein-level IME for downstream analysis. Each set comprises introns detected at all stages of one color and none of the other color. The difference between the median log2 IME scores of the two sets is 0.174, or ∼12%. **D)** Overlaps of iterative FACS-derived sets with RNA-seq-derived sets (DESeq significant introns). See also supplementary Figure S3.

We next explored the relationship between the measurements of RNA and protein: specifically, whether introns in these FACS bins were shifted in their mRNA-level IME estimates as well. Taking the union of all introns seen in a given sort stage (requiring at least 100 reads in either replicate trajectory), we observed that red sorts (R1, R2, R3) had progressively lower and green sorts (G1, G2, G3) had progressively higher mRNA-level IME distributions, indicating consistency between RNA and protein measurements (Fig. 6B). This pattern was independently true for each replicate, and for the intersection of introns detected in both replicates (Fig. S3D). These observations provide orthogonal support for the integrity of our RNA-seq measurements, and for our conclusion that different random introns have distinct effects on gene expression. They further suggest that effects on mRNA levels drive most effects observed at the protein level.

For use in further analysis, we defined a set of “top green introns” as those detected (≥100 reads) in either replicate of all three green sorting stages, and not detected in any red sort; we likewise defined a set of “top red introns” with the opposite requirements. This yielded sets of 676 top green and 462 top red introns, which were well separated in their distributions of RNA-seq-derived IME scores (Fig. 6C). Relatively low overlap was observed with the corresponding significant up or down sets from the DESeq analysis (Fig. 6D), suggesting some unknown source of variability in one or both experimental approaches, at least for introns with more extreme IME values.

### PolyU motifs are enriched in highly enhancing introns

Having sets of relatively stronger and weaker introns in hand, at both the mRNA and protein levels, we sought to identify features enriched in the strongest-enhancing introns. We reasoned that differences in enhancement could be attributable to differences in splicing efficiency, in RNA secondary structure, or in primary sequence, and investigated each possibility accordingly.

First, using a conservative approach that normalizes the GFP:dTom ratio to the proportion spliced, we observed a small but significant positive correlation between splicing efficiency and IME (Fig. 7A, Spearman’s rho=0.072, *p*=2.2e-18; see also Supplemental Note 2): introns with higher proportions of unspliced reads were less likely to be strongly enhancing. However, most introns in the library were efficiently spliced, so other sources of IME variation are clearly present.

**Figure 7.**
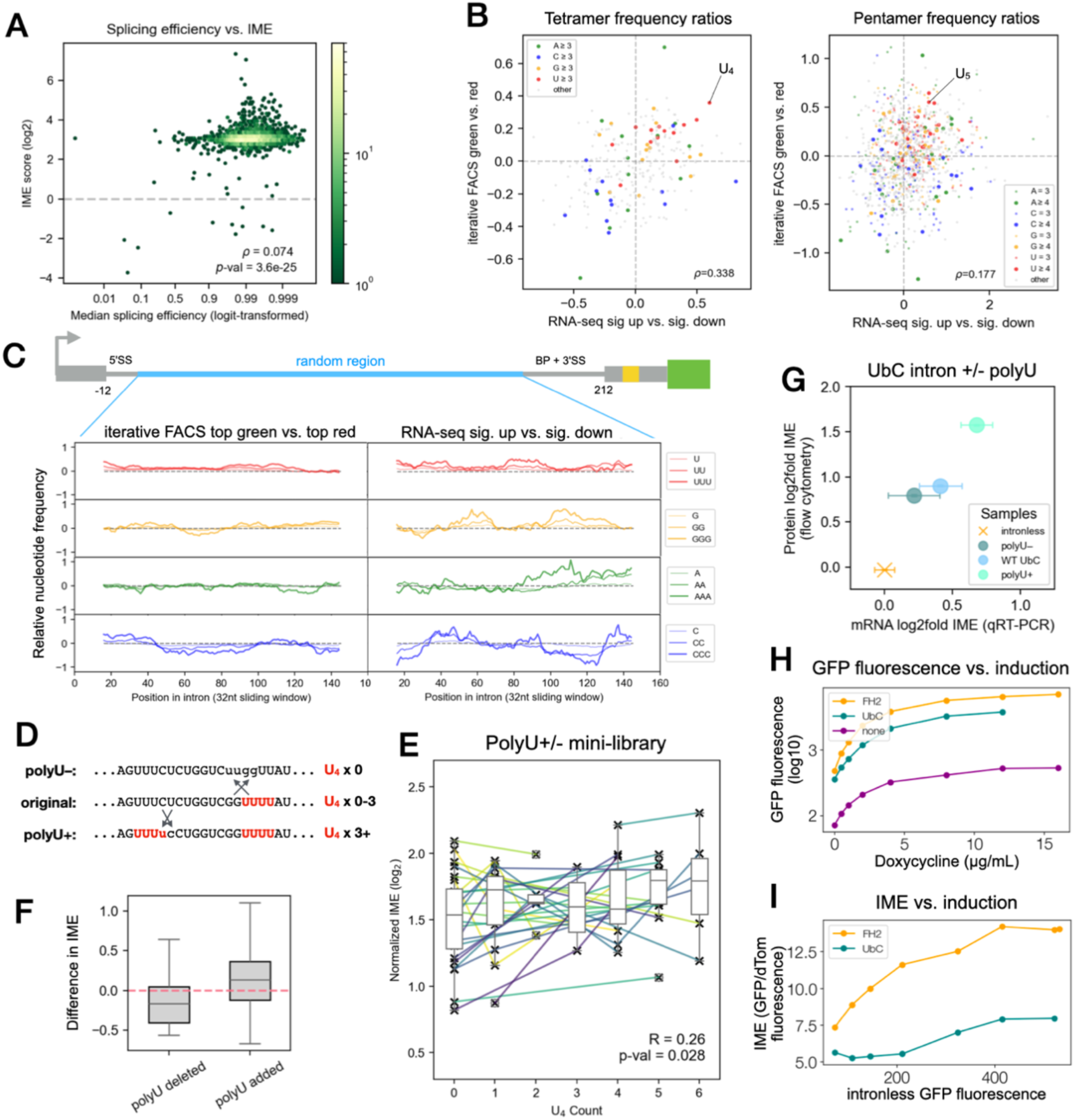
PolyU motifs are enriched in highly-enhancing introns. **A)** Splicing efficiency is weakly but significantly negatively correlated with IME strength. **B)** *K-*mer frequency ratios between highly enhancing and lowly enhancing intron sets for both iterative FACS and RNA-seq. Each dot is coloured according to the nucleotide composition of the *k-*mer and polyU is indicated on both plots (tetramer UUUU, pentamer UUUUU). **C)** Metaplots showing enrichment of single, di- and tri-nucleotide runs in highly vs. lowly enhancing introns, for both iterative FACS and RNA-seq, as a function of position in the intron random region. **D)** Permutation strategy for polyU +/- mini-library. **E)** IME of introns in the second library is significantly correlated with polyU count. Lines indicate permuted versions of the same intron. Boxplot centers represent medians, box edges represent interquartile range (IQR) or middle 50% of data, whiskers extend to 1.5x IQR past each box edge. **F)** Distributions of differences in IME after permutation. **G)** qRT-PCR and flow cytometry of UbC intron polyU series reporters compared to intronless controls. Error bars denote standard error of the mean. **H)** GFP protein expression from a doxycycline-inducible promoter with 5ʹUTR intron indicated in legend. **I)** IME (GFP normalized to dTomato, normalized to intronless) as a function of transcription induction. See also Supplementary Figure S4.

We next explored whether RNA secondary structures were enriched or depleted in introns with higher/lower IME using RNAfold (Hofacker et al., 1994; Lorenz et al., 2011). However, no significant patterns were observed, except for a slight bias against secondary structure downstream of the barcode, in the primer binding region for the RNA-seq amplicon, likely a purely technical effect.

We then interrogated the sequence content of highly versus lowly enhancing introns, considering that short RNA motifs often have biological activity and could conceivably mediate increased IME through, e.g., recruiting *trans*-acting factors to the pre-mRNA. We computed the frequencies of 4-mers and 5-mers in each intron in the library and compared the average frequency of each *k*-mer in the significantly up sets to the corresponding significantly down sets, at mRNA and protein levels (Fig. 7B). We noticed that U-rich *k-*mers were enriched in introns with stronger IME activity, while C-rich *k-*mers tended to be depleted. In particular, polyU motifs (stretches of three or more consecutive Us) were enriched in the introns with the strongest IME from both the FACS and RNA-seq analyses, and the polyU count per intron was slightly but significantly correlated with RNA-seq IME score across the whole library (Fig. S4A, Spearman’s rho = 0.055, *p* = 2e-14, for U_3_).

Comparing the distribution of mRNA-level IME scores between the set of introns lacking polyU and those containing one or more polyU motifs (Fig. S4B), we observed that for all three motif lengths considered (U_3_, U_4_, or U_5_), increased polyU count is generally associated with increased IME (Wilcoxon’s rank-sum test, *p* < 0.05). The differences in the medians of these distributions, taken as a proxy for effect size, are on the order of 0.006-0.018 on a log_2_ scale, or 0.4-1.3% change in observed enhancement. We also broadened this analysis to ask, for all 4-mer and 5-mer motifs, which have significant differences in IME when comparing introns containing them versus introns without them (Fig. S4C). While U-rich *k*-mers again emerged as the strongest differentiators, we also observed that some motifs are associated with a significant decrease in expression – for example the pentamer CGTCA – and that these tended to be either C-rich or A-rich motifs or complex motifs containing three or four distinct nucleotides.

Meta-intron plots of nucleotide enrichment in highly versus lowly enhancing sets also indicate that U-richness is correlated with enhancement (Fig. 7C). In particular, mono-, di- and tri-nucleotide runs of Us are slightly but consistently more frequent across the entire random region of introns with stronger IME, both by iterative FACS and RNA-seq measurements. Though there are local regions of enrichment of other nucleotides – in particular G and A – in the RNA-seq assessment, no base other than U is globally enriched across all positions. Enrichments between iterative FACS bins had generally lower magnitude than those from RNA-seq, perhaps due to differences in the criteria used to define up- and down-regulated sets.

Based on these observations, we designed and queried a second, smaller intron library to directly test the influence of polyU content on IME. We selected 30 random introns from the original library and for each one made “polyU+” and “polyU–” variants, in which we minimally permuted the intron sequence to either bring together or disperse Us, querying 90 sequences in total containing between 0 and 6 instances of U_4_ (Fig. 7D). We sequenced RNA from cells transiently transfected with this pool and found that the IME from these introns was significantly correlated with polyU count (Fig. 7E, Pearson’s R = 0.26, *p* = 0.028). Though neither distribution was significantly different from zero in a one-sample T-test, deleting U_4_s generally led to a slight drop in enhancement, while adding U_4_s led to an increase in enhancement (Fig. 7F).

To further assess the polyU effect in a natural intron context using individual reporter lines, we also created shuffled versions of the human UbC intron sequence. This 812 nt intron natively contains 9 U_4_s (including overlapping motifs, i.e. counting U_5_ as two U_4_s), with our polyU– and polyU+ versions containing 0 and 24 instances of U_4_, respectively. These reporters were integrated into HEK293T cells alongside a matching intronless reporter and observed modest but consistent differences in the IME of these introns (Fig. 7G). The polyU+ version of the UbC intron was stronger in enhancing GFP expression at both the mRNA and the protein level than the original intron, which was in turn stronger than the polyU– intron, confirming that intron polyU content positively modulates IME in a natural sequence context as well as in short random introns.

We noted that the magnitude of fold enhancement versus intronless as well as the baseline expression from this plasmid was lower than for typical introns in the screen, though the plasmid differed in only minor ways from the original backbone (Fig. S4D). Re-cloning these introns into the exact vector used for the screen recovered the magnitude of IME previously observed, but the differences between the polyU+, original and polyU– introns were less pronounced (Fig. S4E). This difference between the vectors suggests that flanking sequence context and/or basal transcriptional output may impact the magnitude of IME and its modulators.

### Other sequence and contextual modifiers of IME

Intrigued by the observation that minor differences in the plasmid backbone altered the IME of introns in our reporter experiments, we decided to explore the effect of promoter strength as a context feature influencing IME. To this end we constructed and integrated one intronless and two intron-containing reporters driven by doxycycline-inducible promoters, and measured the enhancement by each intron compared to the intronless vector at various levels of induction (Fig. 7H). Surprisingly, higher levels of dox induction yielded stronger relative enhancement from both introns (Fig. 7I). Perhaps, in a given genomic context, higher rates of transcription promote more efficient splicing (Bhat et al., 2024; Ding and Elowitz, 2019), which promotes more efficient IME, creating a positive feedback loop between transcription and splicing. This observation also suggests that features of the cellular context (e.g., growth rate, cell cycle phase, etc.) that impact the basal level of transcription of a locus may also impact the level of IME that occurs at that locus, suggesting that IME may act as a magnifier of transcriptionally-driven gene expression programs.

## Discussion

IME has been known for decades, yet our understanding of why different introns can exert different effects in the same context is limited. Here we developed an approach to address this question at an unprecedented scale. Screening tens of thousands of introns with a unique reporter design revealed insights into the sequence determinants of IME, and aspects of the splicing of short introns in human cells.

Our aim was to understand sequence-dependent and -independent contributors to IME. To estimate intron-specific IME levels while accounting for the variability of the data and batch effects, we used DESeq2 to normalize the ratio of GFP to dTom expression per barcode across samples. We found that reporters with introns were expressed on average 8-fold higher than intronless controls, or 5-fold based on qRT-PCR of the pooled library compared to intronless. A likely contributor to this difference is that the RNA-seq mean is the average of per-intron estimates for introns meeting our stringent analysis criteria, whereas the qRT-PCR mean includes many more introns, potentially including a higher proportion of weaker or less efficiently spliced introns. Some individual introns showed stronger or weaker IME than average, suggesting that sequence features can contribute, but these sequence contributions were modest compared to the effect of simply having a well-spliced intron. These intron-specific effects were reproducible in our iterative FACS experiment, further supporting that they reflect properties of particular intron sequences.

Investigating the sequence composition of these introns revealed an enrichment for polyU motifs, and depletion of polyC, in more strongly enhancing introns. PolyU motifs have been reported as enhancing IME in plants (Luehrsen and Walbot, 1994; Rose and Beliakoff, 2000; Clancy and Hannah, 2002; Rose, 2002) but not in animals. In plants, U-richness (or AU-richness) is an important determinant of intron splicing, and depletion of Us can impair splicing. However, deletion studies showed that removal of U-rich motifs can decrease IME without inhibiting splicing (Clancy and Hannah, 2002).

Our data support a functional impact of polyU sequences specifically rather than overall U content, as the polyU reporter experiments involve mutated introns where the nucleotides are shuffled to either disperse Us or gather Us into contiguous stretches, without changing base composition, similar to (Rose et al., 2016). Many families of human RBPs are known to bind short U-rich sequences (Dominguez et al., 2018). Proteins of the PTB family, for example, are known to bind introns and have been reported to enhance mRNA levels by stimulating 3ʹ end processing (Millevoi et al., 2009; Martinson, 2011). It would therefore be interesting to explore whether the IME-promoting activity of polyU or other motifs is dependent on the activities of particular RBPs.

A priori, it was not clear whether introns with 160 nt of random internal sequence would be spliced or not. However, we found that the constant 5ʹ and 3ʹ flanking sequences containing strong SS and a branch point, were sufficient to induce efficient splicing with most internal sequences, as confirmed by RNA-seq and RT-PCR. In rare cases where random introns were not spliced at the expected junction, we could map sites of cryptic splicing internal to the intron. The observed cryptic sites largely matched known 5’ and 3’ SS motifs and obeyed known constraints on minimum intron length in mammals.

Still, the relationship between splicing and IME is complex. Other groups have reported that some IME motifs enhance expression whether located in an intron or exon (Gallegos and Rose, 2019). Transcriptional enhancers can, of course, enhance from outside a gene or from within an intron, where their own transcription may attenuate expression (Cinghu et al., 2017). We observed a small but significant positive correlation between splicing efficiency and IME from random introns, suggesting that the speed of spliceosome assembly or the speed of progression through the spliceosome cycle may contribute to IME.

Curiously, our experiments with an inducible promoter indicated that higher basal levels of transcription from the same locus yields higher IME. Higher transcriptional activity of a locus is known to promote proximity to nuclear speckles, which in turn promotes more efficient splicing (Bhat et al., 2024; Ding and Elowitz, 2019), which we observed to enhance IME. In this context, IME can be considered as one component of a positive feedback loop between transcription and splicing.

Notably, in all our observations of IME, GFP protein levels appeared to be enhanced to a similar degree as GFP mRNA. This suggests that IME acts primarily at the mRNA level in the context studied e.g., via effects on transcription, 3’ end processing, or mRNA stability, with protein levels increased as a result of higher cytoplasmic mRNA levels. Though not explored here, the intron library we have constructed could be used to directly interrogate the impact of each intron on each stage of gene expression, via experiments such as cellular fractionation to assess export, or metabolic labeling to measure rates of mRNA synthesis and stability.

All our experiments, except for the polyU validation library in Fig. 7E, were performed with reporter plasmids integrated at single copy into the genomes of HEK293T cells. This approach was intended to reduce the large variation in plasmid and mRNA levels that occurs with transient transfection, and to study IME in a native chromosomal context. Despite this design, we observed large differences in the magnitude of IME for integration of the same intron between different transfections/integrations of the same plasmid, and for the same intron in distinct but highly similar contexts. These observations suggest that IME is highly sensitive to both the flanking genomic context and to aspects of cell state that may differ between replicate transfection/antibiotic selection regimes, or between the individual cells where plasmids integrated. These properties make it harder to define the IME associated with an individual intron, but suggest that systematic investigation of the impacts of genomic and cellular context on IME might be quite fruitful in uncovering additional contributors to this phenomenon.

## Author Contributions

EJKK conceived the study, designed and performed experiments, analyzed data, and wrote the manuscript. YS performed experiments, analyzed data, and wrote the manuscript. MPM assisted with data analysis. ZJP assisted with reporter experiments. CBB conceived and supervised the study and wrote the manuscript.

## Acknowledgements

We thank Craig Hunter, Sergei Ovchinnikov, Peter Reddien, Phillip Sharp, Seychelle Vos, Dima Ter-Ovanesyan, Conor McMann, Adam Lerer, and members of the Burge lab for helpful discussions. This work was supported by NIH grant 5**-**R01-HG002439 to C.B.B.

## Notes

https://www.ncbi.nlm.nih.gov/geo/query/acc.cgi?acc=GSE278584

https://github.com/ejkk0/IME

### Competing Interest Statement

The authors have declared no competing interest.

