## Supplemental Information for "Sequence determinants of intron-mediated enhancement learned from thousands of random introns"

Supplementary information

Emma J. K. Kowal[1], Yuta Sakai[1], Michael P. McGurk[1], Zoe J. Pasetsky[1], Christopher B. Burge[1,*]

[1]Department of Biology, Massachusetts Institute of Technology, Cambridge MA 02139, USA

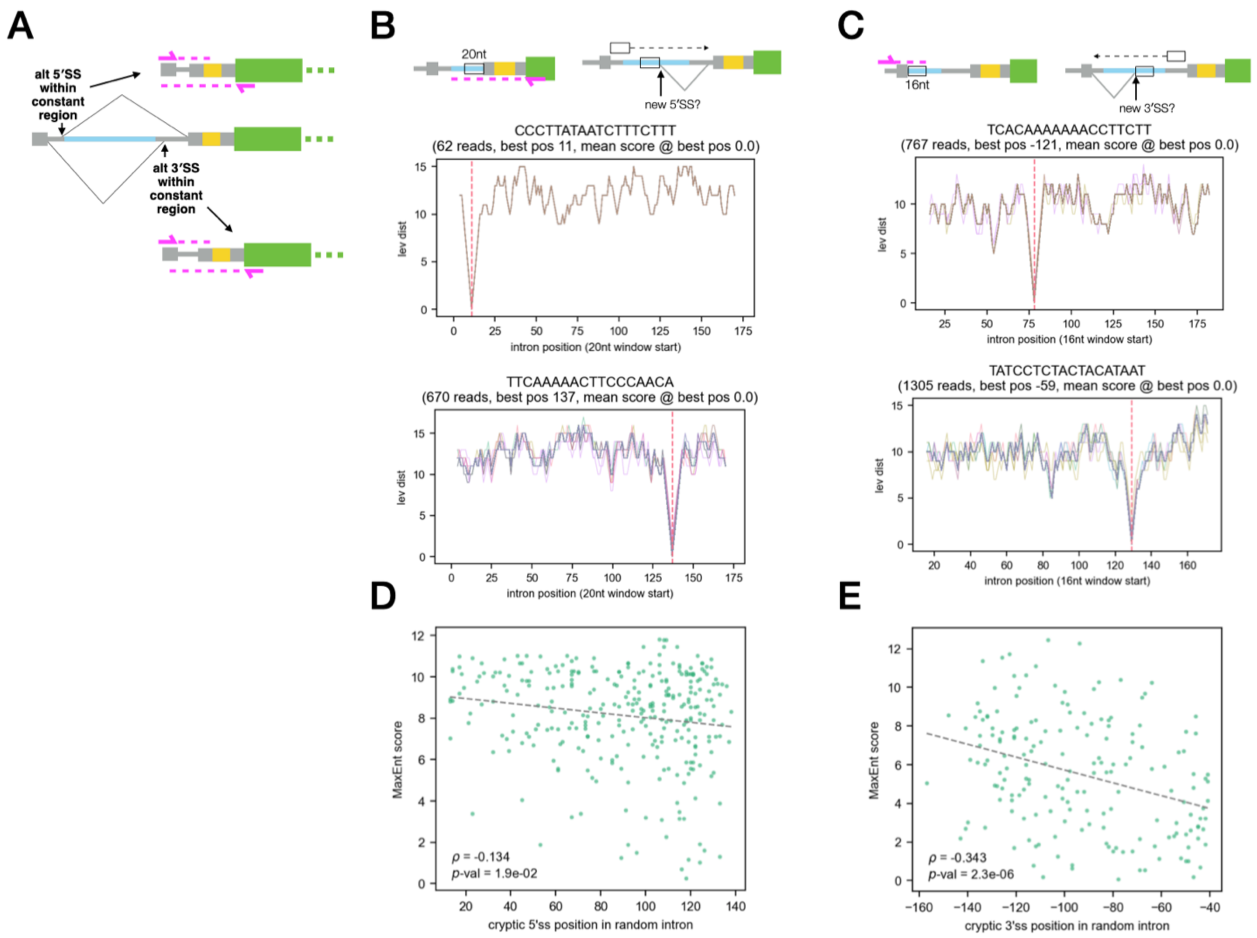


**Figure S1. Identification and analysis of cryptic splice sites (related to Figure 4). A)** Schematic illustration demonstrating how cryptic splicing within the first 28nt or the last 44nt of the intron can alter both ends of the unspliced read. **B)** Schematic of strategy for detecting cryptic 5'SS usage, and example traces showing homology of unspliced read fragments with predicted intron for each read. Introns whose reads all share a single position of best homology, such as those shown, are considered likely substrates of cryptic 5' splicing. **C)** Likewise for cryptic 3'SSs. **D)** Relationship between MaxEnt score of observed cryptic 5'SS and position within the random intron. **E)** As in D, for cryptic 3'SS.

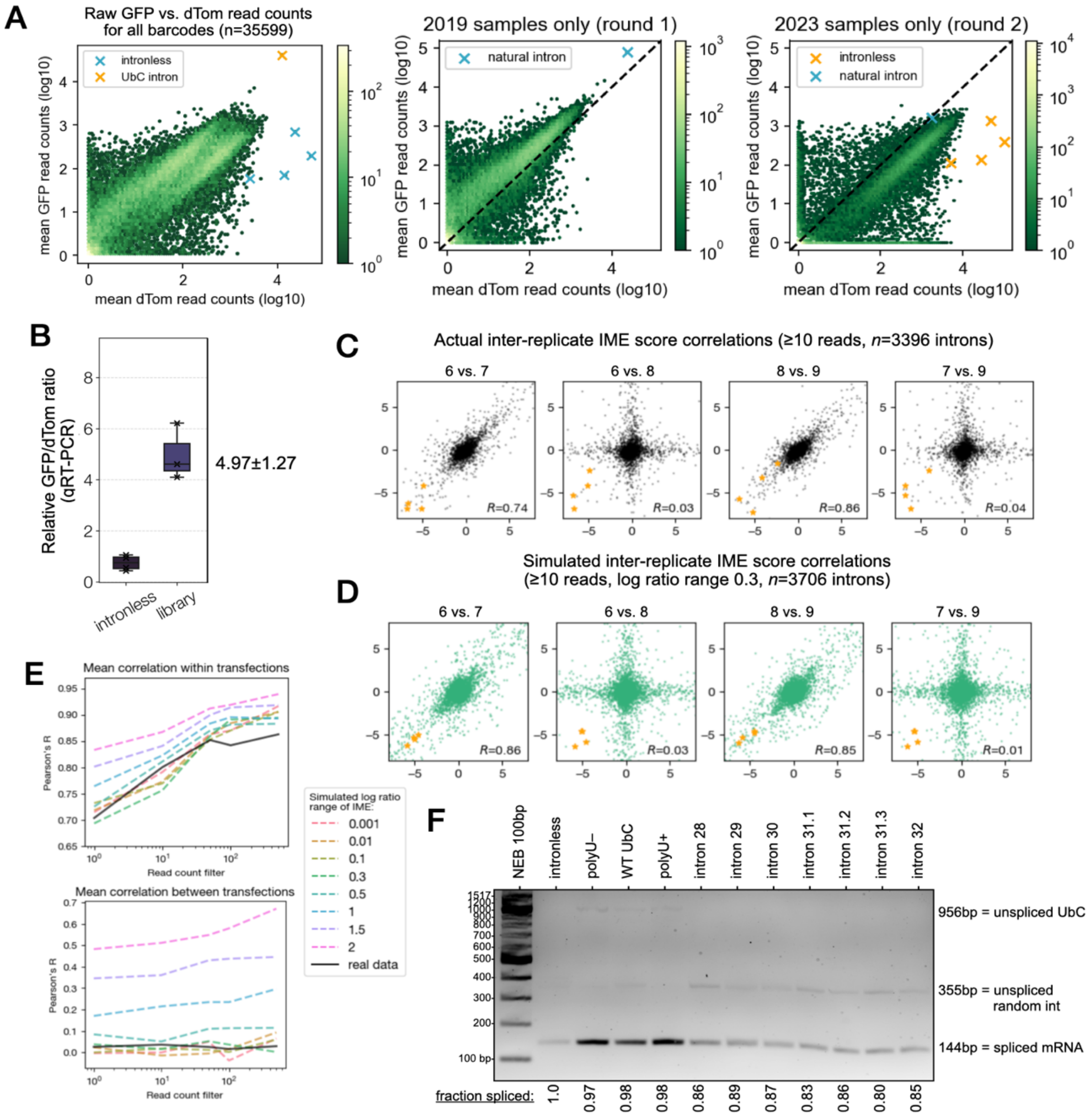


**Figure S2. Estimation and modeling of IME distribution across introns (related to Figure 5). A)** Unfiltered GFP and dTomato read counts for all detected introns across all ten RNA-seq samples. Note how the first five samples have a globally higher GFP/dTom ratio than the last five, performed several years apart. **B)** Bulk qRT-PCR of the random library indicates that the average random intron enhances GFP mRNA expression about five-fold relative to intronless. **C)** Inter-replicate correlations of naive IME scores ($\log_2$ GFP/dTomato read counts) for RNA-seq replicates 6-9, where samples 6&7 originated from one transfection and samples 8&9 from another.. Red stars indicate intronless barcodes. Read count cutoff is enforced for all replicates, not only the two in each plot. **D)** Inter-replicate IME score correlations for simulated read counts based on a model of transfection-independent and transfection-dependent noise. Red stars indicate "intronless" barcodes simulated the same way from a lower starting ratio. **E)** Inter-replicate correlations as a function of read count filter are plotted for real data as well as simulations of different underlying IME strength ranges. **F)** RT-PCR and agarose gel electrophoresis of intron-containing cell lines in Fig. 5E. Expected band sizes for spliced and unspliced amplicons are indicated, as well as relative band intensity of the spliced/spliced+unspliced products for each sample.

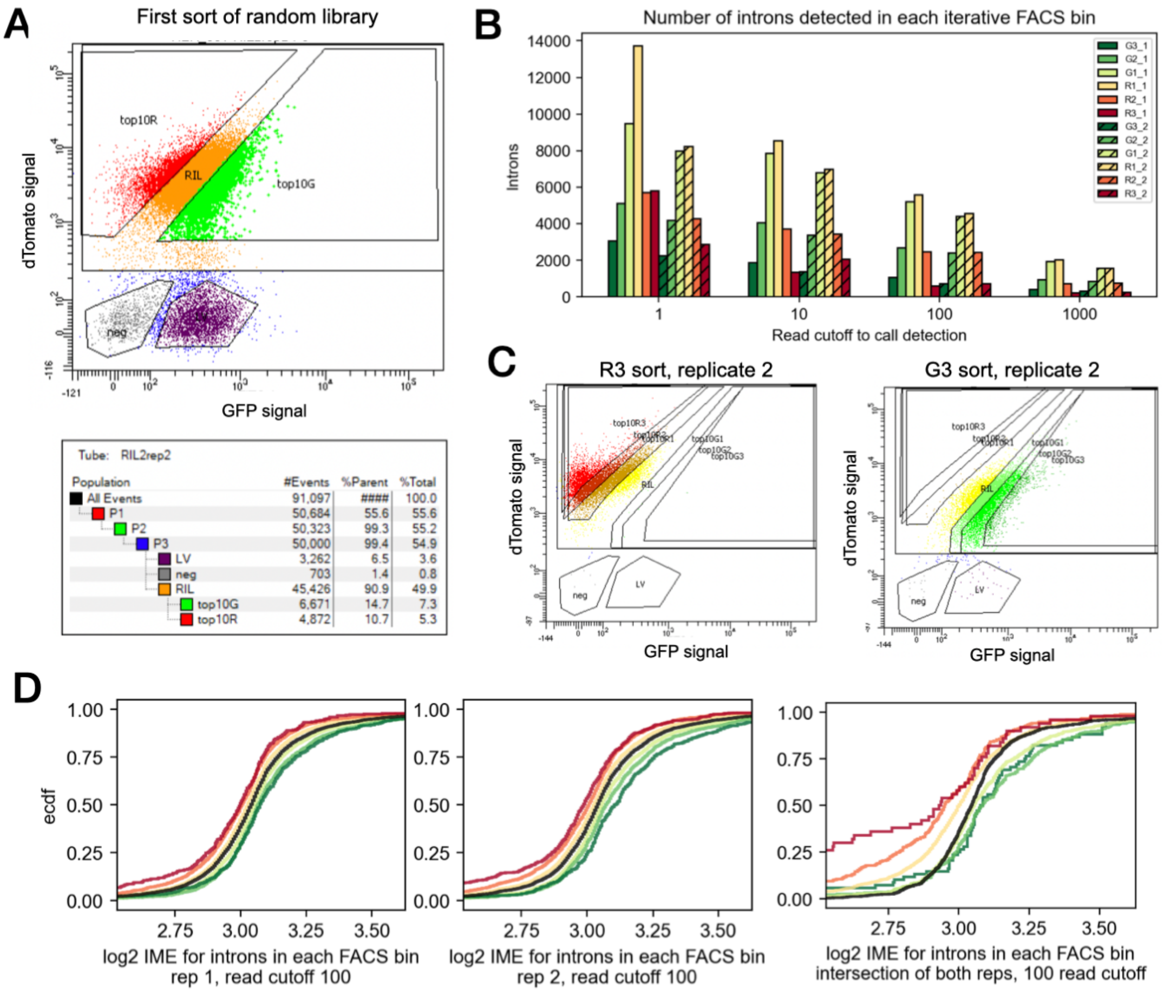


**Figure S3. Iterative selection of introns with high and low IME (related to Figure 6). A)** Raw data from first sort of random library. top10R = R1 sort bin, top10G = G1 sort bin, RIL = Random Intron Library; neg = negative control (parental cell line); LV = library vector, cloning intermediate lacking dTomato. Note that LV composes 6.5% of the overall population. **B)** Number of introns detected in each sort stage of iterative FACS, in both replicate trajectories, as a function of minimum read count for detection. Bars for replicate 2 are hashed. **C)** Representative third stage sorts show that the library mean has shifted significantly from the original population in A. **D)** Both replicate trajectories independently show correspondence with RNA-seq IME scores, as well as the intersection of both.

**A** *k*-mers with significant Spearman correlation count per intron vs. l2fc IME score:

**Positively correlated**

Trimers

| | ρ | *p*-value |
|---|---|---|
| TTT | 0.055 | 2E-14 |
| TCT | 0.034 | 3.04E-06 |
| TTA | 0.028 | 1.24E-04 |
| CTT | 0.027 | 1.92E-04 |
| TTC | 0.026 | 3.01E-04 |
| ATT | 0.026 | 3.07E-04 |
| TCG | 0.026 | 3.28E-04 |
| CTA | 0.025 | 4.27E-04 |

Tetramers

| | ρ | *p*-value |
|---|---|---|
| TTTT | 0.049 | 7.96E-12 |
| CTTT | 0.038 | 1.6E-07 |
| TTTA | 0.037 | 2.83E-07 |
| GTTT | 0.034 | 2.51E-06 |
| TCTT | 0.033 | 5.08E-06 |
| TTTG | 0.031 | 1.5E-05 |
| TTCT | 0.030 | 2.61E-05 |
| TCGT | 0.030 | 3.21E-05 |
| TACT | 0.029 | 6.82E-05 |
| TTAG | 0.028 | 1.02E-04 |
| TTCG | 0.027 | 1.35E-04 |
| TTTC | 0.027 | 1.62E-04 |
| ATTT | 0.027 | 2.03E-04 |
| TAGC | 0.024 | 7.08E-04 |

Pentamers

| | ρ | *p*-value |
|---|---|---|
| ATTTT | 0.033 | 5.62E-06 |
| CTTTT | 0.031 | 1.32E-05 |
| TTTAG | 0.031 | 1.48E-05 |
| TTTTG | 0.030 | 2.51E-05 |
| TCTTT | 0.030 | 3.35E-05 |
| TTTTT | 0.030 | 3.5E-05 |
| GTTTT | 0.028 | 8.16E-05 |
| TTTTA | 0.028 | 1.34E-04 |

**Negatively correlated**

Trimers

| | ρ | *p*-value |
|---|---|---|
| TCA | -0.042 | 4.86E-09 |
| ACA | -0.038 | 1.86E-07 |
| CAA | -0.034 | 2.7E-06 |
| CAC | -0.032 | 1.04E-05 |
| CAG | -0.031 | 1.37E-05 |
| GTC | -0.027 | 1.76E-04 |
| CAT | -0.027 | 2.12E-04 |
| CCA | -0.026 | 2.45E-04 |
| AGG | -0.026 | 3.58E-04 |
| GAC | -0.025 | 6.35E-04 |
| CCC | -0.024 | 7.92E-04 |
| ACG | -0.023 | 1.3E-03 |

Tetramers

| | ρ | *p*-value |
|---|---|---|
| GTCA | -0.041 | 8.52E-09 |
| TGAC | -0.031 | 1.26E-05 |
| ACAA | -0.030 | 2.99E-05 |
| GACG | -0.029 | 6.51E-05 |
| TCAC | -0.028 | 9.5E-05 |
| TCAT | -0.027 | 2.03E-04 |
| GACA | -0.025 | 6.28E-04 |
| CATG | -0.024 | 8.38E-04 |

Pentamers

| | ρ | *p*-value |
|---|---|---|
| CGTCA | -0.035 | 1.13E-06 |
| GTCAA | -0.033 | 3.61E-06 |
| TGACG | -0.033 | 4.98E-06 |
| TGACA | -0.031 | 1.82E-05 |
| TGTCA | -0.030 | 3.26E-05 |
| CGTAA | -0.028 | 1.22E-04 |

**C** *k*-mers with significant Wilcoxon's rank-sum test statistic for introns with 0 vs. ≥1 copy

**Enhancing**

Tetramers

| | effect size | *p*-value |
|---|---|---|
| TTTT | 0.058 | 7.8E-09 |
| CTTT | 0.058 | 6.1E-07 |
| TTTA | 0.054 | 4.9E-06 |
| TTTC | 0.049 | 3.5E-05 |
| TACT | 0.045 | 5.6E-04 |
| TTTG | 0.043 | 1.3E-03 |
| ATTT | 0.042 | 2E-03 |
| TTCG | 0.042 | 2.7E-04 |
| TTAG | 0.040 | 1.5E-04 |
| TCTT | 0.039 | 4.4E-05 |
| TAGC | 0.030 | 9.8E-04 |
| TCGT | 0.030 | 1E-03 |
| GTTT | 0.029 | 1.1E-03 |
| CGGG | 0.023 | 2E-03 |
| TTCT | 0.017 | 3.3E-03 |

Pentamers

| | effect size | *p*-value | | effect size | *p*-value |
|---|---|---|---|---|---|
| AGAAA | 0.056 | 2.2E-03 | GATTT | 0.042 | 8.5E-04 |
| CTACT | 0.052 | 6.9E-04 | TTTGG | 0.041 | 8.1E-04 |
| TCTTT | 0.049 | 6.5E-05 | ATTCG | 0.040 | 1.4E-03 |
| CTTTT | 0.048 | 1.7E-05 | TTTCT | 0.036 | 9.3E-04 |
| TTTAG | 0.045 | 1.5E-05 | TTTTA | 0.035 | 2.9E-04 |
| ATTTT | 0.043 | 3.7E-06 | TTTGT | 0.032 | 2.2E-03 |
| ACTTT | 0.043 | 5.5E-04 | GTTTT | 0.032 | 5.7E-05 |
| TTTTG | 0.043 | 2.7E-05 | TTTTC | 0.031 | 7.4E-04 |
| TTTTT | 0.043 | 7.4E-05 | GGTTT | 0.016 | 1.7E-03 |

**Repressing**

Tetramers

| | effect size | *p*-value |
|---|---|---|
| GTCA | -0.042 | 1E-05 |
| ACAA | -0.041 | 8.4E-05 |
| TCAC | -0.040 | 3E-04 |
| CATG | -0.038 | 2.1E-04 |
| CATC | -0.034 | 3.8E-03 |
| CAAA | -0.032 | 2.2E-03 |
| TCAT | -0.031 | 2.4E-03 |
| ACAC | -0.030 | 2.8E-03 |
| TGAC | -0.029 | 1.6E-04 |
| GACA | -0.025 | 7.2E-04 |
| GACG | -0.024 | 4.4E-04 |

Pentamers

| | effect size | *p*-value | | effect size | *p*-value |
|---|---|---|---|---|---|
| CGTCA | -0.080 | 1E-06 | TGACG | -0.055 | 6.5E-06 |
| CACCC | -0.075 | 2E-04 | GTCAA | -0.055 | 3.9E-06 |
| TACCT | -0.068 | 1.6E-03 | GTCAC | -0.054 | 2.3E-04 |
| ACACC | -0.068 | 9.4E-04 | TCAAA | -0.050 | 2E-03 |
| CAAAC | -0.067 | 5.8E-04 | TGTCA | -0.050 | 4.1E-05 |
| CGTAA | -0.067 | 1.1E-04 | CAGTC | -0.050 | 2E-03 |
| GCAGT | -0.062 | 5.2E-04 | CATCA | -0.049 | 4.5E-04 |
| TGACA | -0.062 | 1.7E-05 | ACAAG | -0.042 | 2.4E-03 |
| GTCAT | -0.055 | 2.8E-04 | GACGT | -0.038 | 1E-03 |
| CATGC | -0.055 | 2.5E-03 | GTCGA | -0.036 | 2E-03 |
| | | | ATATC | -0.034 | 2E-03 |

**B**

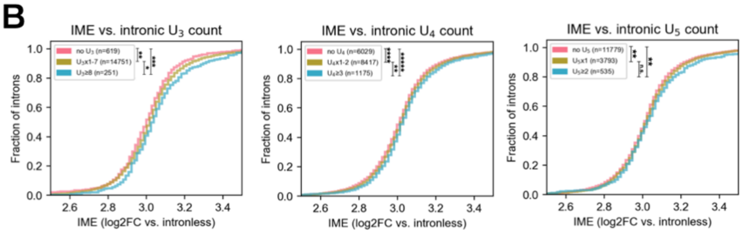


**E**

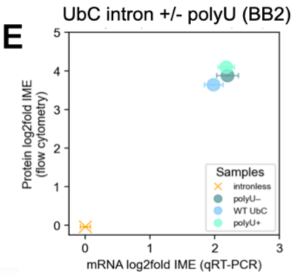


**D**

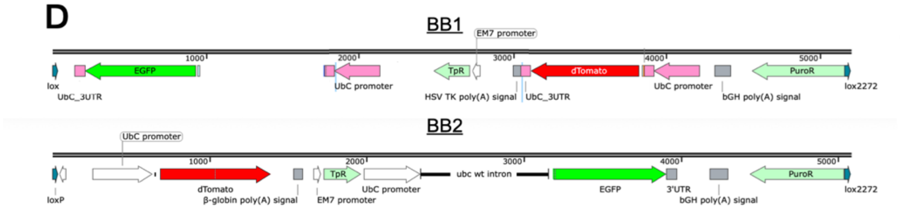


**Figure S4. Sequence motifs associated with stronger and weaker IME (related to Figure 7). A)** Splicing efficiency is weakly but significantly negatively correlated with IME strength. Grey dashed line indicates intronless IME. **B)** *k*-mer frequency ratios between highly enhancing and lowly enhancing intron sets for both iterative FACS and RNA-seq. Each dot is coloured according to the nucleotide composition of the *k*-mer. PolyU is indicated on both plots (tetramer UUUU, pentamer UUUUU). **C)** Metaplots showing enrichment of single, di- and tri-nucleotide runs in highly vs. lowly enhancing introns, for both iterative FACS and RNA-seq, as a function of position in the intron random region. **D)** Experimental design of polyU +/- series library and reporter experiment. **E)** A smaller second library of 30 random introns plus shuffled versions increasing or decreasing polyU content (90 introns total) show a positive correlation between polyU count and IME. Boxplot centers represent medians, box edges represent interquartile range (IQR) or middle 50% of data, whiskers extend to 1.5x IQR past each box edge. **F)** qRT-PCR and flow cytometry of polyU series reporters compared to intronless controls. Error bars denote standard error of the mean. **G)** GFP protein expression from a doxycycline-inducible promoter with or without 5'UTR intron. Legend indicates intron present in construct. **H)** IME (GFP normalized to dTomato, normalized to intronless) as a function of transcription induction. Legend indicates intron present in construct. See also supplementary Figure S4.

### Supplemental note 1: Simulation

In later replicates of the RNA-seq experiment, parallel transfections of the library were performed and cells were labelled to indicate which original transfection had produced them. This design decision confirmed something we had suspected, that there is a significant source of transfection-specific noise in our experimental workflow. We conceived the simulation shown in figure S2 as an attempt to understand the curious shape of the scatter plots of IME between inter-transfection versus intra-transfection replicates. The simulation architecture which successfully reproduced these shapes suggests the presence of multiple sources and types of noise: large perturbations to read counts could be created by random differences in efficiency of cell proliferation or PCR amplification, both of which are exponential processes in which small initial variation can lead to large differences in outcome.

We hypothesize that occasional extreme IME values which were shared within transfections and not across them represent some relatively rare but consequential events during transfection, integration, and selection; for example, off-target integration, transcriptional silencing of GFP or dTomato, or perhaps simply large variation in transfection efficiencies combined with imperfect removal of barcode mismatches. These events could also help to explain the lack of correspondence between predicted IME strengths of individual introns from bulk measurements and validation of IME strength from individual transgenic lines as in figure 5E.

Off-target integration was assessed in the original paper describing development the HILO-RMCE system, and found to happen at a rate of ~0.04 copies per diploid genome in the HeLa A12 cell line (Fig. S3, Khandelia et al., 2011). This is roughly compatible with our simulated addition of high-variance transfection-specific read count noise to one in ten introns. It should also be noted, importantly, that the enrichment of higher and lower RNA-seq IME scores by iterative FACS (Fig. 6B) cannot be entirely explained by this transfection-specific noise, as the CDFs of IME score by FACS bin still separate correctly if bins derived from one transfection are used with IME scores computed only using the opposite transfection (data not shown).

### Supplemental note 2: IME and splicing

It is important to emphasize that our estimates of IME are normalized to the proportion of transcripts that are spliced, since we are interested here in the impact of splicing on gene expression. Consider an intron spliced in only 10% of transcripts that has the same high ratio of spliced GFP / dTomato reads as an intron spliced in 100% of GFP transcripts. The poorly spliced intron achieves the same boost in functional, mature mRNA production with only 10% as many splicing events, and thus *per splicing event*, its IME potency is 10X that of the well-spliced intron. To fairly compare these introns, we must divide the raw IME score of the first intron by 0.1 (its splicing efficiency). Thus, the splicing-efficiency-corrected IME score is defined as the ratio of GFP/dTomato reads, normalized to the intronless ratio, and then normalized to the splicing efficiency of the intron (ratio of spliced to total GFP reads).

In this way, our metric emphasizes the impact of the splicing of an intron on the production of spliced transcripts from its locus. The measure is conservative in the sense that, if unspliced transcripts are less stable than spliced transcripts, splicing efficiency will be over-estimated, tending to underestimate IME. Thus, our conclusion that low splicing efficiency is associated with lower IME occurs despite the conservativeness of our IME measure. It is also possible that our amplicon sequencing compresses the range of splicing efficiencies present, which might also tend to underestimate the magnitude of this effect.
